# Relating the past with the present: Information integration and segregation during ongoing narrative processing

**DOI:** 10.1101/2020.01.16.908731

**Authors:** Claire H. C. Chang, Christina Lazaridi, Yaara Yeshurun, Kenneth A. Norman, Uri Hasson

## Abstract

This study examined how the brain dynamically updates event representations by integrating new information over multiple minutes while segregating irrelevant input. A professional writer custom-designed a narrative with two independent storylines, interleaving across minute-long segments (ABAB). In the last (C) part, characters from the two storylines meet and their shared history is revealed. Part C is designed to induce the spontaneous recall of past events, upon the recurrence of narrative motifs from A/B, and to shed new light on them. Our fMRI results showed storyline-specific neural patterns, which were reinstated (i.e. became more active) during storyline transitions. This effect increased along the processing timescale hierarchy, peaking in the default mode network. Similarly, the neural reinstatement of motifs was found during part C. Furthermore, participants showing stronger motif reinstatement performed better in integrating A/B and C events, demonstrating the role of memory reactivation in information integration over intervening irrelevant events.

## Introduction

Real-life events unfold over multiple minutes. Using real-life stimuli such as stories and movies, previous studies have revealed a cortical hierarchy of timescales that synthesize information over increasing temporal receptive windows (TRWs) (Baldassano et al., 2017; J. Chen et al., 2016; Honey, Thesen, et al., 2012; Lerner, Honey, Silbert, & Hasson, 2011; Yeshurun, Nguyen, & Hasson, 2017). To address these findings, we proposed a process-memory model (Hasson, Chen, & Honey, 2015). Unlike classic theories of working memory, which distinguish between areas that process incoming information and working memory buffers that accumulate and protect the processed information (Baddeley, 2003; Cowan, 2008), the process-memory model posits that all cortical areas actively sustain memories while dynamically synthesizing them with newly arrived input at their preferred timescales. Namely, early sensory areas integrate information over short timescales of tens of milliseconds, coinciding with the duration of phonemes and words. Adjacent areas along the superior temporal cortex integrate information over hundreds of milliseconds, coinciding with the duration of single sentences, while high order areas, which overlap with the default mode network (DMN) (Buckner, Andrews-Hanna, & Schacter, 2008; Raichle et al., 2001), integrate information across paragraphs as the narrative unfolds over many minutes. This framework illustrates a simple recurrent mechanism for continuous event updating in long processing timescale areas at the top of the hierarchy. However, in real life, we often have to integrate discontinuous pieces of information to develop a full understanding of an event. This raises the question of how areas with long processing timescales integrate incoming information with relevant past events, while, at the same time, preserving the accumulated information and protecting it from being integrated with irrelevant current events.

To probe this question, we collaborated with a professional author (C.L) to craft an original fictional story with a purposefully designed narrative structure. The first part of the narrative consisted of two seemingly unrelated storylines, A, which takes place in Los Angeles, employing one set of characters, and B, which takes place in New York and involves a distinct set of characters (Figure 1). The two storylines were presented in an interleaved fashion over 30 segments, 15 segments for each storyline (A1B1A2B2…A15B15). In the last 15 segments (Part C), characters from the two storylines meet in New York and their shared history is revealed. In other words, the C part updates the two storylines with new information previously unknown to the audience. One of the main techniques for bridging part C with A and B was to embed specific images/situations/phrases, i.e., narrative motifs, within either the A or B storylines. The recurrence of these motifs in part C was designed to reinstate specific moments from storylines A and B. For example, in segment A1, the main character, Clara, makes homemade chili for her husband in LA. In part C, Clara eats and comments on a B storyline character’s (Steven’s) homemade chili recipe, which is designed to reactivate the memory of segment A1 and update it by revealing that Clara was making chili following Steven’s example. To develop a full understanding of this story, the listeners need to piece together a series of clues (motifs and storylines) like a detective.

**Figure 1.**
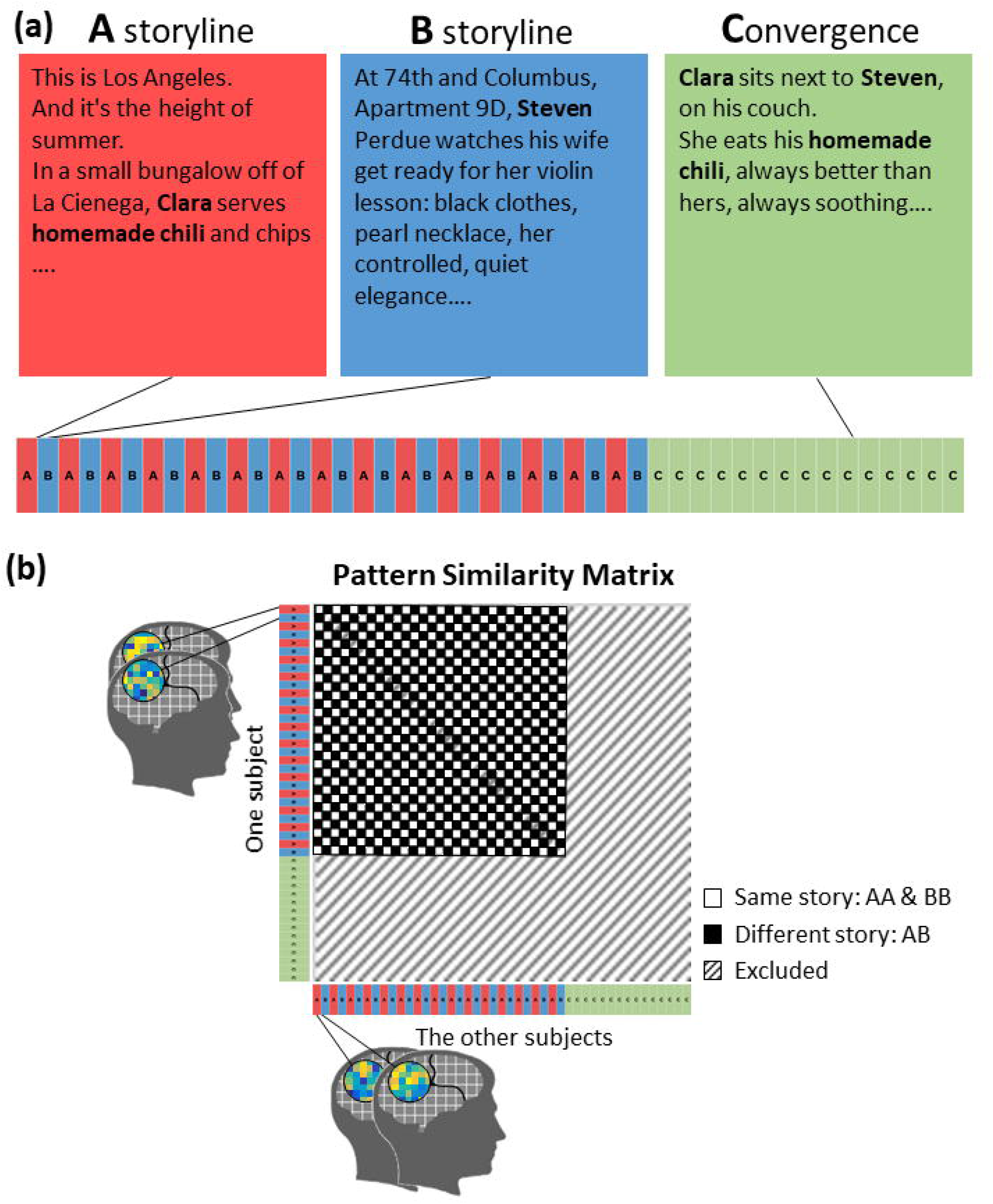
Stimulus and design. (a) The stimulus is a spoken narrative composed of 45 segments with two storylines, A and B. A and B interleave in the first thirty segments and converge in the last fifteen segments (C part). (b) Predicted similarity matrix between brain responses to the 45 segments.

We hypothesized that, within individual A/B segments, high-level cortical regions would integrate information over time (i.e., they would show process memory) in order to build up a representation of the ongoing situation. In keeping with Event Segmentation Theory (Zacks et al., 2001; Zacks & Swallow, 2007), we further hypothesized that storyline switches would result in “flushing out” of the previous storyline’s representation in high-level cortical regions (see also Chien & Honey, 2020 & Ezzyat & Davachi, 2011), making it possible for that region to start representing the features of the other storyline. We also hypothesized that episodic memory would play a key role in reinstating existing storyline representations when the narrative returned to those storylines.

These hypotheses led to the following predictions about the A/B period of the story: First, we predicted that high-level cortical regions would contain distinct neural representations for the A and B storylines, as the storylines depict different situations and (according to Event Segmentation Theory) these distinct situation models will be flushed out at storyline switches, minimizing “carryover” of the neural patterns. Second, we predicted that hippocampus would be engaged at storyline switches in order to reactivate stored episodic memories of the relevant storyline and that the degree of hippocampal engagement would predict the degree of activation of the relevant storyline representation; we measured hippocampal engagement using hippocampal-cortical inter-subject functional correlation (ISFC; Simony et al., 2016). In a previous study (J. Chen et al., 2016), we observed stimulus-driven ISFC between hippocampus and DMN cortical regions when the participants resumed a movie after a 1-day break; here, we tested whether hippocampal-cortical connectivity would also facilitate the reinstatement of recent memories from minutes ago as listeners switch from one storyline to the other. Lastly, we predicted that neural representations of the A and B storylines would become more distinct over the course of the narrative, reflecting the accumulation (across segments) of distinctive details pertaining to each segment.

During part C, we hypothesized that the recurrence of motifs from the A/B storylines (e.g., the chili recipe) in part C would induce spontaneous recall of related past events and that recalling these A/B events (in response to a motif) would help participants to understand the relationship between the A/B segments and the narrative in part C. To test this, we quantified the “reminding” effect during part C by measuring whether motifs triggered neural reinstatement of corresponding moments in A/B events. To look at how this affected behavior, we computed a “relation score” (using a post-scan test) that specifically tracked participants’ understanding of how the motifs connected A/B events with the narrative in part C. We predicted that, across participants, the degree of neural A/B reinstatement triggered by motifs would be correlated with these behavioral relation scores.

Our study is not the first to examine the interleaving of storyline representations and the recurrence of narrative motifs with fMRI. In one relevant study, Milivojevic et al. (2016) used an audio-visual movie with interleaved storylines (“Sliding Doors”) and found coding of storyline information in the hippocampus (but not in the DMN areas that are the focus of our study). In another relevant study, Kauttonen et al. (2018) scanned participants who watched the movie “Memento”, in which recurring cues were embedded to trigger memory recall, similar to the motifs in our story; they found that recurring scenes activated matching, scene-specific neural patterns in low-level sensory regions, and also that there was a common (i.e., not-scene-specific) neural pattern that was evoked in DMN regions whenever participants viewed a scene for a second time.

An important difference between our study and these other studies is that they used pre-existing narrative movies as stimuli, whereas we used a stimulus that was custom-crafted to address the hypotheses enumerated above. Our use of a custom-crafted stimulus allowed us to build on these prior studies while avoiding confounds that were present in these other studies. A limitation of the design of Milivojevic et al. (2016) is that storyline differences were confounded with sensory differences (e.g., related to some locations being more prevalent in one storyline than the other). Our study resolves this confound because both storylines were presented auditorily by the same speaker and thus did not differ in low-level sensory properties. Likewise, a limitation of the design of Kauttonen et al. (2018) is that recurring scenes were repeated exactly; as such, pattern similarity between matching cues could be due to matching sensory inputs (as opposed to memory). In contrast, in our study, recurring motifs were embedded in different scenes, making it easier to distinguish between the influences of perception and memory. By eliminating these confounds while still providing an engaging stimulus, we hoped to provide a more detailed view of how incoming information is segregated from irrelevant memories and integrated with relevant memories as a narrative unfolds over time.

## Methods

### Participants

Twenty-eight participants were recruited. They were all right-handed native English speakers. All participants provided written consent forms before the experiment. Twenty-five participants were included for further analyses (14 females, age 18-40). Three were excluded, one due to anatomical anomalies, one due to excessive motion artifacts in T1 image, and one slept during the story. The experimental protocol was approved by the Institutional Review Board of Princeton University.

### Stimulus

The stimulus was created by Lazaridi (“The 21st Year” -- Excerpt, copyright 2019), who has been in collaboration with our lab for a number of years (Yeshurun, Swanson, et al., 2017). She has years of experience in practicing and evolving the technique of organizing the audience’s understanding, memory, and interpretation of a narrative through screenplay writing and professional screenplay development around the world (Lazaridi, 2012). Compared to other types of writing, the creation of a screenplay is highly audience-driven due to the large investment (in time, collaboration, and financing) inherent in film-making. Furthermore, watching a film is a more continuous experience than reading a book, requiring the screenwriter to guide and unite the audience’s understanding and overall response to the narrative without loss of focus or inner thought digressions.

Lazaridi designed the narrative stimulus as a stand-alone fiction text that incorporated her experience-guided narrative techniques of traditional screenplay writing. The narrative consisted of 45 segments, and two seemingly unrelated storylines, A and B. A and B segments were presented in an interleaved manner for the first 30 segments. In the last 15 segments (Part C), the two storylines merged into a unified narrative. Each segment lasted for 41-57 TRs (mean: 46 TRs = 70 sec). They were separated by silent pauses of 3-4 TRs. The narrative was recorded by a professional actress (June Stein), who is a native English speaker, and directed by Lazaridi to ensure that the actor’s interpretation matched the author’s intent. The recording is 56 minutes long (see Supplementary Table 1 for the transcription of the story). In the A and B segments, the author incorporated unique narrative motifs, i.e., specific images/situations/phrases that recurred in part C (see Figure 1 and Figure 5 for a sample motif and Supplementary Table 2 for a list of all the motifs). The recurrence of motifs in part C is designed to trigger the reinstatement of specific moments from part AB, in order to evolve their meanings with new information from the C part. The same narrative motif was not always realized with the same words (“throwing up” vs. “Clara feels sick, as the coffeecake rises to her throat”). In total, there were 28 different narrative motifs, occurring 58 times in the AB part, and 36 times in part C.

### Procedure

The recording of the narrative was presented using MATLAB 2010 (MathWorks) and Psychtoolbox 3 (Brainard, 1997) through MRI-compatible insert earphones (Sensimetrics, Model S14). MRI-safe passive noise-canceling headphones were placed over the earbuds for noise reduction and safety. To remove the initial signal drift and the common response to stimulus onset, the narrative was preceded by a 14 TR long musical stimulus, which was unrelated to the narrative and excluded from fMRI analysis. Participants filled a questionnaire after the scanning, to evaluate their overall comprehension of the narrative and their ability to relate events in different parts of the story that shared the same motifs.

### MRI acquisition

Subjects were scanned in a 3T full-body MRI scanner (Skyra, Siemens) with a 20-channel head coil. For functional scans, images were acquired using a T2*-weighted echo planar imaging (EPI) pulse sequence (repetition time (TR), 1500 ms; echo time (TE), 28 ms; flip angle, 64°), each volume comprising 27 slices of 4 mm thickness with 0 mm gap; slice acquisition order was interleaved. In-plane resolution was 3 × 3 mm^2^ (field of view (FOV), 192 × 192 mm^2^). Anatomical images were acquired using a T1-weighted magnetization-prepared rapid-acquisition gradient echo (MPRAGE) pulse sequence (TR, 2300 ms; TE, 3.08 ms; flip angle 9°; 0.86 × 0.86 × 0.9 mm^3^ resolution; FOV, 220 × 220 mm^2^). To minimize head movement, subjects’ heads were stabilized with foam padding.

### MRI analysis

#### Preprocessing

MRI data were preprocessed using FSL 5.0 (http://fsl.fmrib.ox.ac.uk/) and NeuroPipe (https://github.com/ntblab/neuropipe), including BET brain extraction, slice time correction, motion correction, high-pass filtering (140 s cutoff), and spatial smoothing (FWHM 6 mm). All data were aligned to standard 3 mm MNI space (MNI152). Only voxels covered by all participants’ image acquisition area were included for further analysis.

Following preprocessing, the first 19 TRs were cropped to remove the music preceding the narrative (14 TRs), the time gap between scanning and narrative onset (2 TRs), and to correct for the hemodynamic delay (3 TRs). To verify the temporal alignment between the fMRI data and the stimulus, we computed the temporal correlation between the audio envelop of the stimulus (volume) and the subjects’ mean brain activation in left Heschl’s gyrus following Honey et al. (2012). The left Heschl’s gyrus mask was from Harvard-Oxford cortical structural probabilistic atlases (thresholded at 25%). The audio envelope was calculated using a Hilbert transform and down-sampled to the 1.5 s TR. The correlations were computed with −100-100 TRs lag to find the time lag that showed the highest correlation. The averaged peak time was 0.12 TR across subjects, indicating that the narrative and fMRI data were temporally well-aligned.

To account for the low-level properties of the stimulus, a multiple-regression model was built for each voxel. The regressors included an intercept, the audio envelope, and the boxcar function of the between-segment pauses, convolved by the canonical hemodynamic response function and its derivatives with respect to time and dispersion as given in SPM8 (https://www.fil.ion.ucl.ac.uk/spm/). For the effect of audio amplitude and between-segment pause, please see Supplementary Figure 1. The residuals of the regression model were used for the following analyses.

#### ROI masks

We used 238 functional regions of interest (ROIs) defined independently by Shen et al. (2013) based on whole-brain parcellation of resting-state fMRI data. A control anatomical ROI was also included: left Heschl’s gyrus defined using the Harvard-Oxford cortical structural probabilistic atlas, thresholded at 25%.

A bilateral anatomical hippocampal mask was obtained using the same threshold. It has been proposed that the temporal integration window of episodic memory representation varies along hippocampal long axis (Collin, Milivojevic, & Doeller, 2015). While the posterior hippocampus has a small representation scale, the middle and anterior portions are able to contain the associations between more than two events. If so, it would be inappropriate to treat hippocampus as a functionally homogenous region. Therefore, we divided the hippocampus mask into anterior (MNI coordinate y > −19), middle (−30 < y <= −19), and posterior parts (y <=−30) ROIs following Collin et al. (2015).

All ROIs had more than 50 voxels in our data.

#### Shared response model

When comparing activation patterns across subjects, the mismatch of functional topographies could decrease analysis sensitivity even after anatomical alignment (Brett, Johnsrude, & Owen, 2002; Sabuncu et al., 2010). Therefore, we functionally aligned data within each ROI across subjects using the shared response model (SRM) (Brain Imaging Analysis Kit, http://brainiak.org) (P.-H. Chen et al., 2015). SRM projects all subjects’ data into a common low-dimensional feature space by capturing the components of the response shared across subjects. The input to SRM was a TR x voxel x subject matrix, and the output was a TR x feature x subject matrix. We used fMRI data from the whole story (z-scored over time first) to estimate an SRM with 50 features. Note that no information about storyline or motif was submitted to SRM. Therefore, while this projection inflated the overall inter-subject pattern similarity, it could not artifactually give rise to the storyline or motif effect shown here. The output of SRM was z-scored over time. Unless otherwise stated, all the pattern analyses described below were run based on the resulting 50 features.

We also performed the same analyses without the application of SRM. Generally speaking, a subset of the areas that were significant in the analysis with SRM were also significant in the analysis without SRM. Please see Supplementary Figure 2 for the results.

#### RSA of storyline effect

To examine the storyline effect, we performed representational similarity analysis (RSA) (Kriegeskorte, Goebel, & Bandettini, 2006; Kriegeskorte, Mur, & Bandettini, 2008) on brain activation patterns and tested whether the representational similarity between segments from the same storyline was higher than that of segments from different storylines. We first computed the averaged activation within each segment across TRs for each voxel. The resulting 45 values were then z-scored across segments. For each ROI, pairwise pattern similarities between the 45 activation maps were computed with the leave-one-subject-out method (Figure 1b). Namely, the averaged activation pattern was extracted for each segment. Then the Pearson correlation coefficients between one subject’s activation patterns and the averaged patterns of the remaining subjects were computed. The output correlation coefficients (45 x 45 segments) were normalized with Fisher’s z-transformation. This procedure was repeated for each of the 25 subjects and each ROI.

We then contrasted the averaged within- and between-storyline similarities in the AB part, excluding the within-segment similarities (the diagonal of the 45 x 45 similarity matrix), to obtain 25 contrast values (Figure 2a) for each ROI. These contrast values were compared to zero by a one-tailed one-sample t-test and thresholded at p < .05 (FWE correction for multiple comparisons). The results were projected back onto the whole-brain surface and visualized using Freesurfer v6 (http://surfer.nmr.mgh.harvard.edu/).

**Figure 2.**
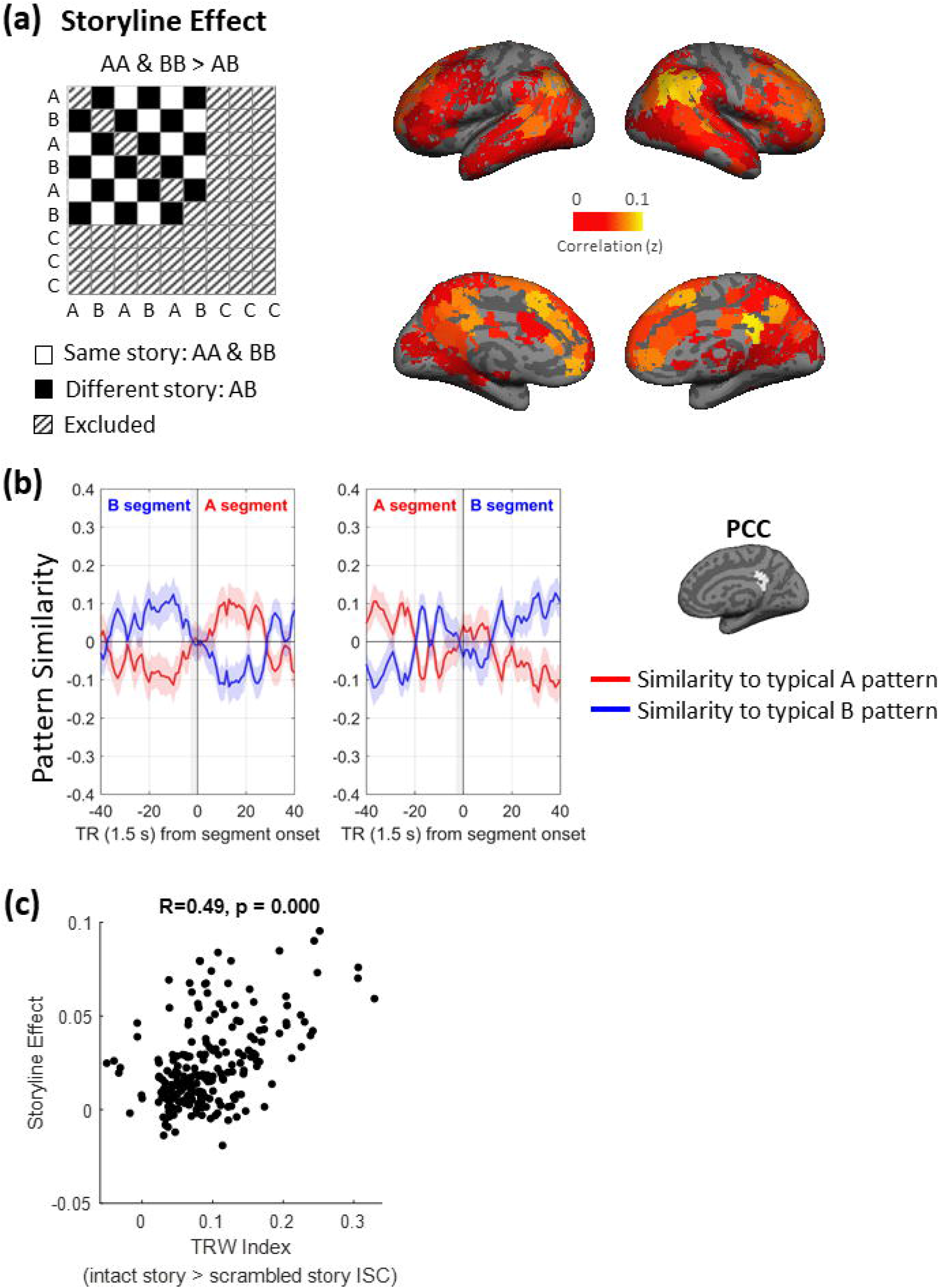
Storyline effect. (a) Regions showing larger pattern similarities between segments of the same vs. different storylines (N=25, p < .05, FWE). (b) Pattern similarity between typical A or B storyline patterns and −40~40 TRs surrounding the segment boundaries in right PCC. Shaded areas indicate 95% confidence interval (CI) across subjects. The vertical gray-shaded area shows the silent pause at boundary. (c) Correlation between temporal receptive window index and storyline effect across regions.

To examine whether the storyline effect increased over time, for regions showing a significant storyline effect, we computed the storyline effect in the early (segment 1-14) and later (segment 15-30) halves of the A/B part separately. 25 contrast values were generated by comparing the late and early storyline effects (late (same > different storyline) > early (same > different storyline)). These contrast values were again submitted to a one-tailed one-sample t-test (p < .05, FWE) (Figure 3a).

**Figure 3.**
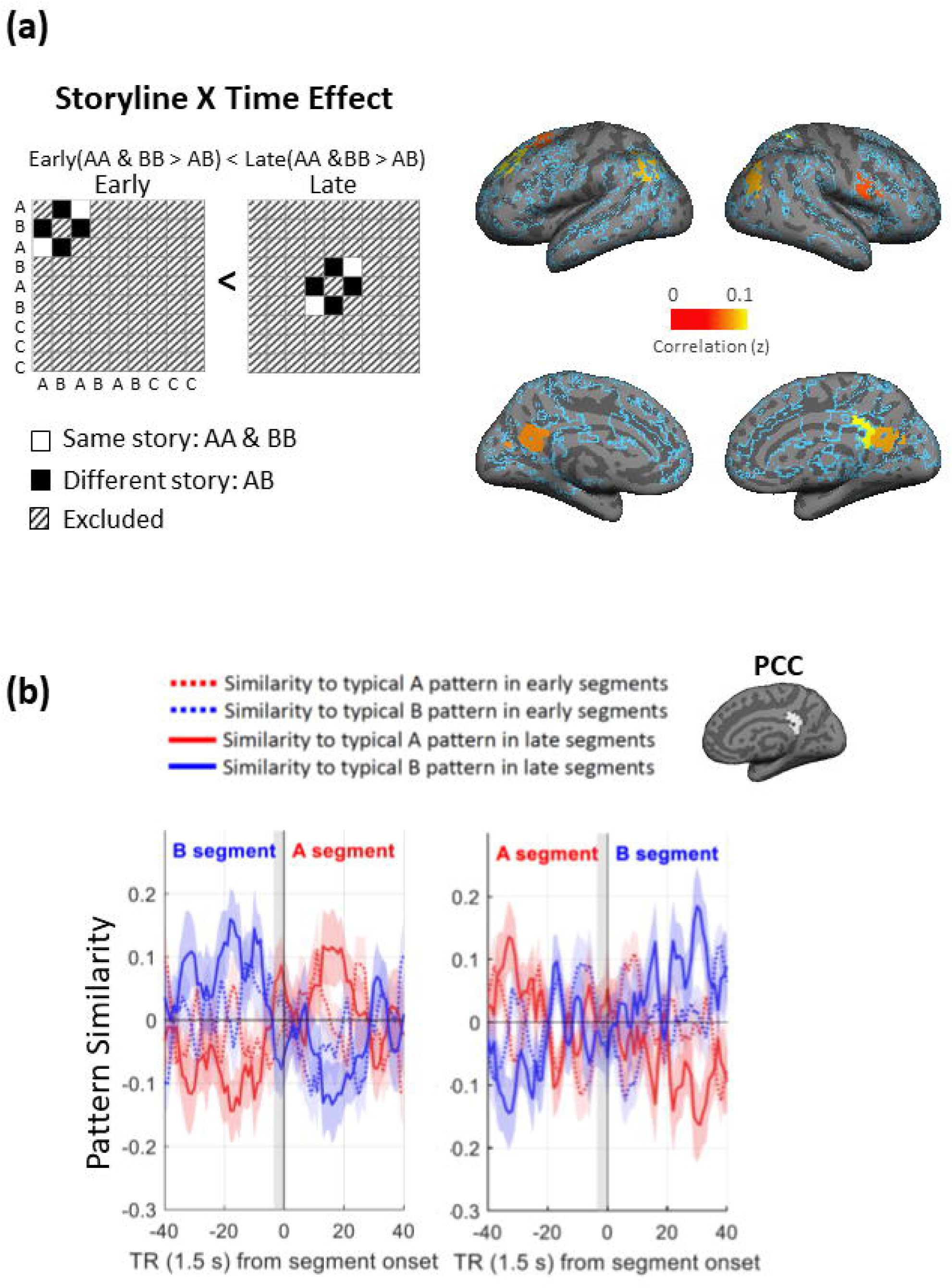
Storyline x Time effect within regions showing significant storyline effect. (a) Regions showing larger storyline effect in the late than early half of the AB part (N=25, p < .05, FWE). The blue outline marks regions showing a significant storyline effect. (b) Pattern similarity between typical A or B storyline patterns and −40~40 TRs surrounding segment boundaries in right PCC, computed for the early and segments separately. Shaded areas indicate 95% CI across subjects. The vertical gray-shaded area shows the silent pause at boundary.

To test the storyline x time effect in a more graded way, we constructed a 45 x 45 time effect matrix, populated with the average of the time points (segment number). For example, the (4, 5) entry of this matrix is 4.5 (=(4+5)/2). Taking only the entries corresponding to within-storyline similarity, excluding the diagonal elements (Supplementary Figure 3, upper panel), we computed the Pearson correlation between the time matrix and the pattern similarity matrix. The resulting R-values were entered into a one-sample one-tailed group t-test after Fisher’s z-transformation within regions showing significant storyline effect (N=25, p < .05, FWE). The between-storyline dissimilarity was tested separately in a similar manner. The overlap between these two effects was shown in the lower panel of Supplementary Figure 3.

#### Time course of the storyline effect at segment boundary

To further illustrate the time course of the storyline effect, we computed the pattern similarities between each of the −40-40 TRs around segment onsets and the typical A and B storyline patterns using a leave-one-subject-out method. For example, for the boundary between segment 1 and segment 2, −40~40 TRs around the onset of segment 2 were extracted from one subject. The typical A storyline pattern was obtained by averaging all the A storyline TRs, except for the segments analyzed here, i.e. segment 1 and 2, from the rest of the subjects. The typical B storyline pattern was obtained in the same manner. Pearson correlation between the 81 TRs around segment 2 onset and the typical A and B patterns were calculated and normalized with Fisher’s z-transformation. The same procedure was repeated for each subject and each boundary. Please see Figure 2b for the results.

To further illustrate the time x storyline effect, we applied the above analysis to the early and late segments separately. Namely, typical A and B patterns were computed using early and late segments respectively. TRs around the boundaries between early segments were compared with the early templates and TRs around the boundary between the late segments were compared with the late templates. The results are shown in Figure 3b.

#### Temporal receptive window index

Following Yeshurun et al. (2017), the TRW index was generated based on an independent dataset from Lerner et al. (2011), which includes an intact story (“Pieman”, ~7 min long) and the same story with scrambled word order. Inter-subject correlation between averaged time series of each ROI was computed, using the leave-one-subject-out method, and normalized using Fisher’s z transformation. TRW index was then calculated by subtracting the ISC of the scrambled story from that of the intact story. We examined the correlation between TRW and storyline effect across regions (Figure 2c).

#### RSA of narrative motif effect

For each narrative motif occurrence, we obtained the corresponding activation pattern by averaging 5 TRs immediately after its onset based on the intuition that motif effect was transient and lasted only for a few sentences. Pearson correlation coefficients between activation patterns of motifs in the AB part and motifs in the C part were computed with the leave-one-subject-out method and normalized with Fisher’s z transformation. As shown in Figure 5, pattern similarities between narrative motifs were grouped into three types: (1) same motif, (2) different motifs from the same storyline, and (3) different motifs from different storylines (unrelated). For example, pattern similarities between different occurrences of “chili” belong to (1). Similarities between “chili” and other A storyline motifs belong to (2). Similarities between chili and B storyline motifs belong to (3). Motif effect of each “chili” token in part C was defined as the averaged type (1) similarity minus the averaged type (2) similarity, in order to eliminate the confound of storyline effect.

The group motif effect was thresholded with a permutation test. For each ROI, the above procedure was repeated after shuffling the labels of motifs within storylines for 10000 times, creating a null distribution. To correct for multiple comparisons across ROIs, the largest motif effect across ROIs in each of the 10000 iterations was extracted, resulting in a null distribution of the maximum motif effects. Only ROIs with a group motif effect exceeding 95% of the null distribution were considered significant (p < .05, FWE) (Figure 6, middle).

#### Motif-related events with shared vs. different main characters

To verify that the motif effect does not only reflect shared characters, the author of the story designated the main character in each motif-related event and a predicted pattern similarity matrix was generated based on whether two events shared the same character. We ran a permutation test similar to the one described above and found no significant character effect (p < .05, FWE) (Supplementary Figure 4).

#### Time course of the narrative motif effect

To further illustrate the time course of the motif effect, for each motif in C, the Pearson correlation coefficients between activation patterns of −5~10 TRs around its onset and the activation patterns of motifs in the AB part were computed. Motifs with a −5~10 TRs time window that overlapped with the between-segment silent pauses were excluded from this analysis. The resulting coefficients were normalized with Fisher’s z-transformation and averaged by categories (same motif and same storyline, different motif but same storyline, and unrelated). For each ROI, we applied two-tailed paired t-tests to compare pattern similarities between categories at each time point (p < .05, FWE correction for time points). Figure 6 shows the resulting pattern similarity around narrative motif onset.

#### Narrative motif vs. high-frequency word effect

To verify that the motif effect did not result from repeated wordings or word-level semantics, we replaced the narrative motifs with storyline-specific high-frequency words and performed the same RSA. More specifically, among words that only occurred in A and C parts and words that occurred only in B and C parts, we chose the 28 words with the highest lemma/word stem frequencies (Supplementary Table 3). Two out of the twenty-eight narrative motifs were included in this list. Together, these words occurred 111 times in the AB part and 110 times in the C part. Among regions showing a significant motif effect, we calculated the difference between the real motif effect and the effect elicited by high-frequency words for each subject. The 25 difference values were entered into a one-sample one-tailed t-test. The results were thresholded at p < .05 (FWE, Figure 7).

#### Within-subject RSA of the storyline and motif effect

Based on our prior work showing that between-subject analysis is able to reveal the shared coding of events across subjects (Baldassano et al., 2017; Baldassano, Hasson, & Norman, 2018; J. Chen et al., 2017; Zadbood, Chen, Leong, Norman, & Hasson, 2017) and that it also boosts signal-to-noise ratio (Simony et al., 2016), we adopted between-subject RSA as our “default” in this study; however, we also included within-subject RSA for comparison purposes.

The results of our within-subject RSA analyses are shown in Supplementary Figures 5-8. Considering the potential impact of temporal autocorrelation (Mumford, Davis, & Poldrack, 2014) and low-frequency drift (Alink, Walther, Krugliak, van den Bosch, & Kriegeskorte, 2015) in the fMRI signals on within-subject similarity matrix, especially between neighboring segments, we also included storyline analyses thresholded using the label permutation method (Supplementary Figure 9). For the storyline effect, we shuffled the labels of segments 1-30 10000 times to obtain a null distribution of the group mean effect. This procedure was performed for each ROI and the resulting p-value was corrected for multiple comparisons across ROIs (FWE). The storyline x time effect was tested within regions that showed a significant storyline effect by shuffling segment labels within storylines.

#### Correlation between hippocampal-cortical ISFC and cortical reinstatement of storyline and motif

To examine whether the cortical reinstatement of storyline was dependent on connectivity with the hippocampus, we examined the Pearson correlation between hippocampal-cortical inter-subject connectivity (ISFC) (Simony et al., 2016) and the storyline effect across segments for each subject, within ROIs showing a significant storyline effect.

The ISFC was computed within the 0-40 TR time window after the onset of each segment using the leave-one-subject-out method, i.e. the correlation between one subject’s hippocampal activity and the averaged cortical activity of the other subjects. We used the preprocessed data without regressing out the effects of between-segment pause and the audio envelope because it is possible that the activation pulse between segments (Supplementary Figure 1) does not only reflect the silence but also memory encoding or retrieval (Ben-Yakov & Dudai, 2011). SRM was not applied because topographical alignment is not a concern when comparing the averaged time series between ROIs. The hippocampus seed was defined using Harvard-Oxford cortical structural probabilistic atlas thresholded at 25%.

The storyline effect for each segment was also computed using the leave-one-subject-out method. The typical activation patterns for A and B storylines were first estimated by averaging data from all but one subject, excluding the current segment. Pattern similarity between the resulting typical A & B patterns and the left-out subject’s activation pattern for the current segment was then computed. The storyline effect was defined as the difference between pattern similarity to the relevant storyline and the similarity to the irrelevant storyline, taking the previous segment as the baseline, for example, for a B segment: (current segment’s similarity to B - similarity to A) - (previous segment’s similarity to B - similarity to A).

The correlation between ISFC and the storyline effect across segments was computed for each subject, excluding the first segment of each storyline. The R-values were entered into a one-tailed t-test after Fisher’s z-transformation. This initial analysis did not yield a significant result after correction for multiple ROIs (N = 25, p < .05, FDR correction). In exploratory follow-up analyses, we then systematically examined the influence of the time window of ISFC, the time window of the storyline effect, the hippocampus seed (whole vs. posterior: MNI y <= −30), and the baseline of the storyline effect. We also examined the correlation across subjects in each segment. For each combination of analysis parameters, we corrected for multiple-comparisons across ROIs using the FDR method. Please see Figure 9 for the results.

We examined the correlation between hippocampal ISFC and motif reinstatement in a similar manner. The motif effect was defined and computed using the RSA method described above (based on 5 TRs after motif onsets, using similarity between different motifs from the same storylines as a baseline). Across motifs in the C part, the correlation between the motif effect and ISFC after motif onset was then computed for each subject. We also examined the correlation across subjects for each motif and the influence of ISFC time windows and hippocampus seeds.

## Results

fMRI data were collected from 25 subjects while they listened to a structured narrative that lasted for approximately 1 hour. The narrative has two interleaved, seemingly unrelated storylines, A & B, that converge in the later C part. In the first set of analyses, we tested how ongoing information from each of the two unrelated storylines was accumulated across minute-long segments while being segregated from the parallel unrelated interleaved storyline. In the second set of analyses, we tested how events in part A & B were reactivated in part C. The two storylines are connected to part C using 28 specifically-designed, recurrent narrative motifs. These motifs were planted at specific, strategic moments of the narrative by the author (58 occurrences in parts A & B, and 36 occurrences in part C). Participants’ understanding of the relations created by these motifs was assessed based on post-scan questionnaires.

For the fMRI data, we first regressed out the effect of audio amplitude and between-segment pause (Supplementary Figure 1) and applied the shared response model (SRM) to adjust for the mismatch of functional topographies across subjects (P.-H. Chen et al., 2015)(please see Supplementary Figure 2 for results without applying SRM). Using representational similarity analysis (RSA) (Kriegeskorte et al., 2006, 2008) on brain activation patterns within ROIs independently defined by a whole-brain parcellation of resting-state fMRI (Shen et al., 2013), we tested whether the structure of the story induced the reinstatement of storylines and narrative motifs and whether this led to the integration of separate events.

### Neural reinstatement of storyline

We first examined whether, and if so, where in the brain the two seemingly unrelated storylines (A & B) had distinct cortical representations. Using RSA, we compared the neural patterns within each storyline (AA & BB) to the neural patterns between the two storylines (AB). Within each ROI, we averaged over time within each segment (lasting approximately one minute) to extract a spatial pattern of activity for that segment. We then compared pattern similarity between segments from the same storyline to pattern similarity between segments from different storylines (Figure 2a).

Higher within-storyline pattern similarity was revealed in a large set of regions, including language areas (superior/middle temporal gyrus, inferior frontal gyrus, and supplementary motor cortex), areas in the default mode network (including posterior cingulate gyrus, precuneus, medial prefrontal cortex, superior frontal gyrus, posterior parietal cortex, angular gyrus, posterior hippocampus, and parahippocampal cortex), areas in the executive network, (including anterior insula, middle temporal gyrus, middle cingulate gyrus, and supramarginal gyrus), high order visual areas (including cuneus and fusiform gyrus), and subcortical areas (including putamen, thalamus, and caudate).

Figure 2b shows the time course of storyline effect at the segment boundary in the region where the largest separation across the two storylines was found, i.e. posterior cingulate gyrus (PCC)/precuneus. We computed the pattern similarity between each of the −40~40 TRs around segment boundaries and the typical A or B storyline patterns. At the boundary between B and A segments, the similarity to the typical B pattern rapidly dropped, while the similarity to the typical A pattern increased. The two waveforms crossed around the boundary. Similar results were obtained for the complementary transition from A to B segments.

It is worth noticing that, although the two curves in Figure 2b seem to be symmetrical with respect to zero, that does not mean that the two storylines had opposing activation patterns. The two patterns are forced to average to approximately zero by the need to subtract the global mean response before computing the typical A/B patterns (Garrido, Vaziri-Pashkam, Nakayama, & Wilmer, 2013; Murphy, Birn, Handwerker, Jones, & Bandettini, 2009). Therefore, the correlation values only reflect the relative, but not the absolute, difference between storylines.

### Stronger neural reinstatement of storyline in areas with longer processing timescales

We used an independent dataset (Lerner et al., 2011) to generate a temporal receptive window (TRW) index for each ROI, i.e. the difference in inter-subject correlation between an intact story and its scrambled version. Higher TRW indices were found in prior studies to be associated with increased capacity to accumulate information over long-timescales (Lerner et al., 2011; Yeshurun, Nguyen, et al., 2017). If the storyline effect only reflected a difference in low-level properties such as wording or acoustic features (note that the same narrator read all segments), regions with low TRW, i.e. regions insensitive to word scrambling, should also show a storyline effect as strong as in high TRW regions. On the contrary, we found a significant positive correlation between TRW index and storyline effect (Figure 2c). In other words, areas that are capable of accumulating information over long-timescale had a larger difference between storylines.

### Storyline x Time effect

We predicted that the segregation of the two storylines (A & B) should increase as the story unfolds and subjects accumulate further information about the unique context of each storyline. To test this hypothesis, we examined whether the storyline effect increased over time by dividing the AB part into the early and later halves (Figure 3a). Within areas showing the separation between storylines, an increase in the separation of patterns at the later phase (leading to a significant interaction between time and storyline) was found in PCC/precuneus, left angular gyrus/inferior parietal lobule, left superior frontal gyrus, right inferior frontal gyrus, right middle temporal gyrus/middle occipital gyrus, and right superior parietal lobule. Figure 3b shows the storyline transition at segment boundaries in the early and late AB part respectively.

We compared the storyline effect in the early and late time bins to avoid imposing assumptions on the time effect (e.g. linearity). Having said this, a graded time x storyline effect is observed in similar regions (Supplementary Figure 3).

We also examined pattern similarity in anatomically defined hippocampus ROIs and again observed the separation between storylines (Figure 4). The 3-way interaction between subregions (anterior/middle/posterior), time (early/late), and storyline (same/different) is significant (F(2, 48) = 20.84; p < .001). Post-hoc two-tailed paired t-tests showed a same > different storyline effect in the late AB part in anterior hippocampus (t(1,24) = 2.07, p = .048, Cohen’s d = 0.41, CI = 0.0001~0.041), in the early AB part in the middle hippocampus (t(1,24) = 5.91, p <.001, Cohen’s d = 1.18, CI = 0.03~0.07), and both the early (t(1,24) = 3.47, p = .002, Cohen’s d = 0.69, CI = 0.01~0.05) and late AB part (t(1,24) = 5.25, p < .001, Cohen’s d = 1.05, CI = 0.03~0.07) in the posterior hippocampus.

**Figure 4.**
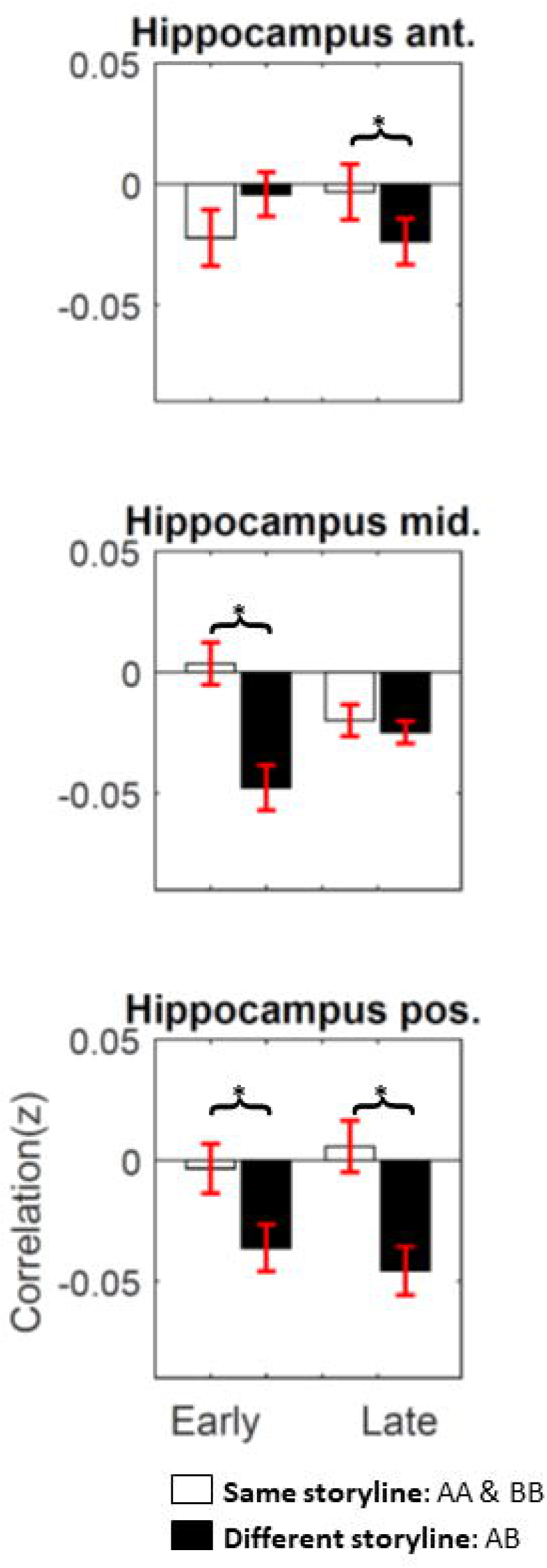
Storyline effect in hippocampus. Error bars indicate 95% CI across subjects. The 3-way interaction between subregions (anterior/middle/posterior hippocampus), time (early/late), and storyline (same/different) is significant (p < .001). The asterisks indicate significant post-hoc two-tailed paired t-test (p < .05).

### Neural reinstatement of narrative motifs

Information in the C part sheds new light on both A and B events, e.g. Clara learned the chili recipe from Steven; Margaret’s mustard-stained blouse now reminds Steven of her death. We examined how past information from part AB was reinstated during part C upon the recurrence of the narrative motifs. For each occurrence of motifs in the story, we averaged the 5 TRs after its onset. Then, we correlated each reoccurrence of a narrative motif in part C with all its occurrences in part A or B (Figure 5). The correlation between matching motifs was computed, as well as the correlation between non-matching motifs from the same storyline (shared storyline) and the correlation between non-matching motifs from the competing storyline (unrelated segments).

**Figure 5.**
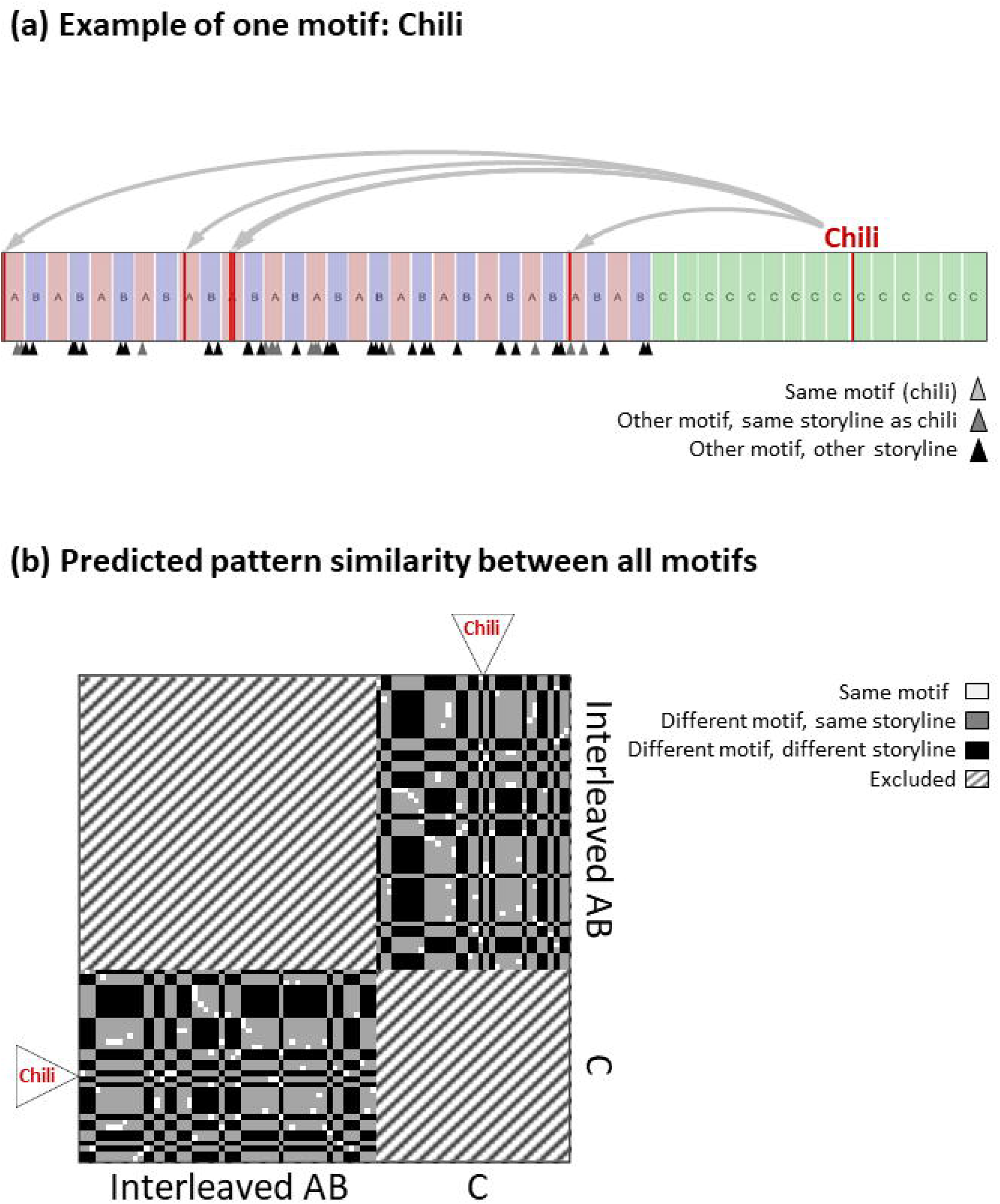
Predicted motif effect. (a) Events from the convergence part (C part) were bridged to specific events in the AB part by motifs, e.g. chili. (b) Predictions about pattern similarity based on motifs.

Compared to non-matching motifs from the same storyline, the reappearance of the narrative motifs in part C reinstated specific neural patterns seen when the motifs were encountered during the A/B segments in PCC/precuneus, bilateral clusters in posterior temporal lobe/inferior parietal lobes/higher visual areas, bilateral lateral frontal areas, and dorsal medial prefrontal cortex (dmPFC) (Figure 6).

**Figure 6.**
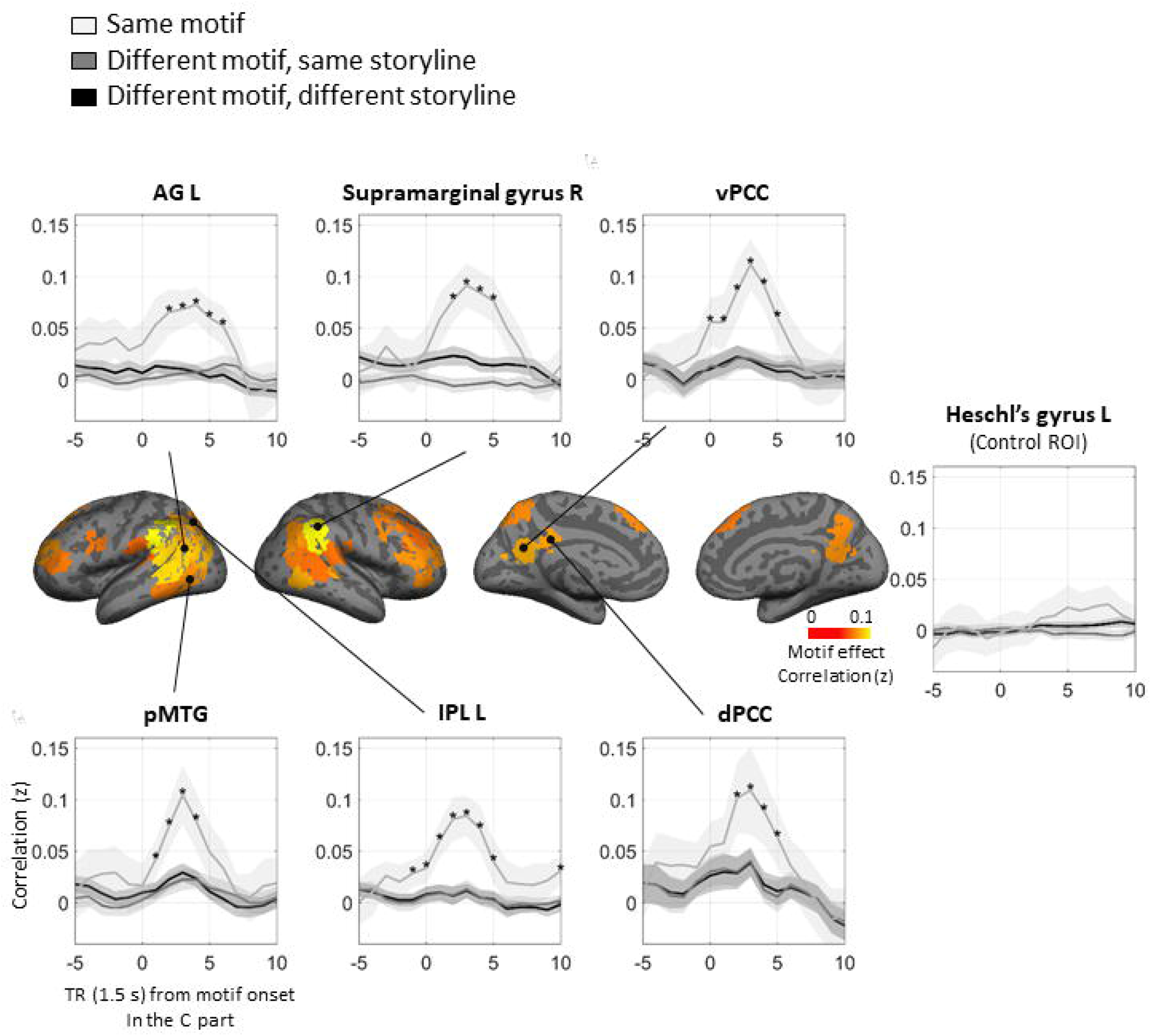
Motif effect. Middle: regions showing a significant motif effect (p < .05, FWE). Upper & Lower: Pattern similarities between motifs in the AB part and TRs around motif onsets in the C part. Shaded areas indicate 95% CI across participants. Asterisks mark time points showing larger similarity between the same motifs than both control categories (p < .05, FWE correction for time points).

Furthermore, TR by TR analysis around the onsets of narrative motifs in part C showed that the correlation rapidly increased after motif onset and lasted for 4-7 TRs, approximately 3-6 sentences (Figure 6, upper and lower panels). The reinstatement effect was specific to matching motifs and was not seen between non-matching motifs, either within or across storylines.

To verify that the motif effect does not only reflect shared characters, we compared entries with shared vs. different main characters in the motif x motif pattern similarity matrix and found no significant character effect (Supplementary Figure 4).

### Narrative motifs vs. high-frequency word effects

To make sure that the reinstatement of patterns after motif onsets reflects the retrieval of narrative information (as opposed to simple reactivation of word representations shared between the A/B and C segments, e.g., the representation of the word “chili”), we performed the same analysis on a set of high-frequency words that occurred in the C part and in either the A or B storyline (e.g., “watch”). We analyzed 28 high-frequency words to match the number of narrative motifs. If the neural reinstatement effect that we observed for motifs simply reflected the reactivation of word representations, the same effect should be observed when we look at the repetition of high-frequency words like “watch” that have no particular narrative significance. In all ROIs showing a significant motif effect, besides the dorsal PCC, the correlation between matching items was significantly higher for the narrative motifs compared to the high-frequency words, for which the correlation hovered around zero (p < .05, FWE corrected, Figure 7). This indicates that word repetition alone was not sufficient to drive the motif reinstatement effect we observed; rather, the words had to refer to significant narrative events (as is true for “chili” but not for “watch”).

**Figure 7.**
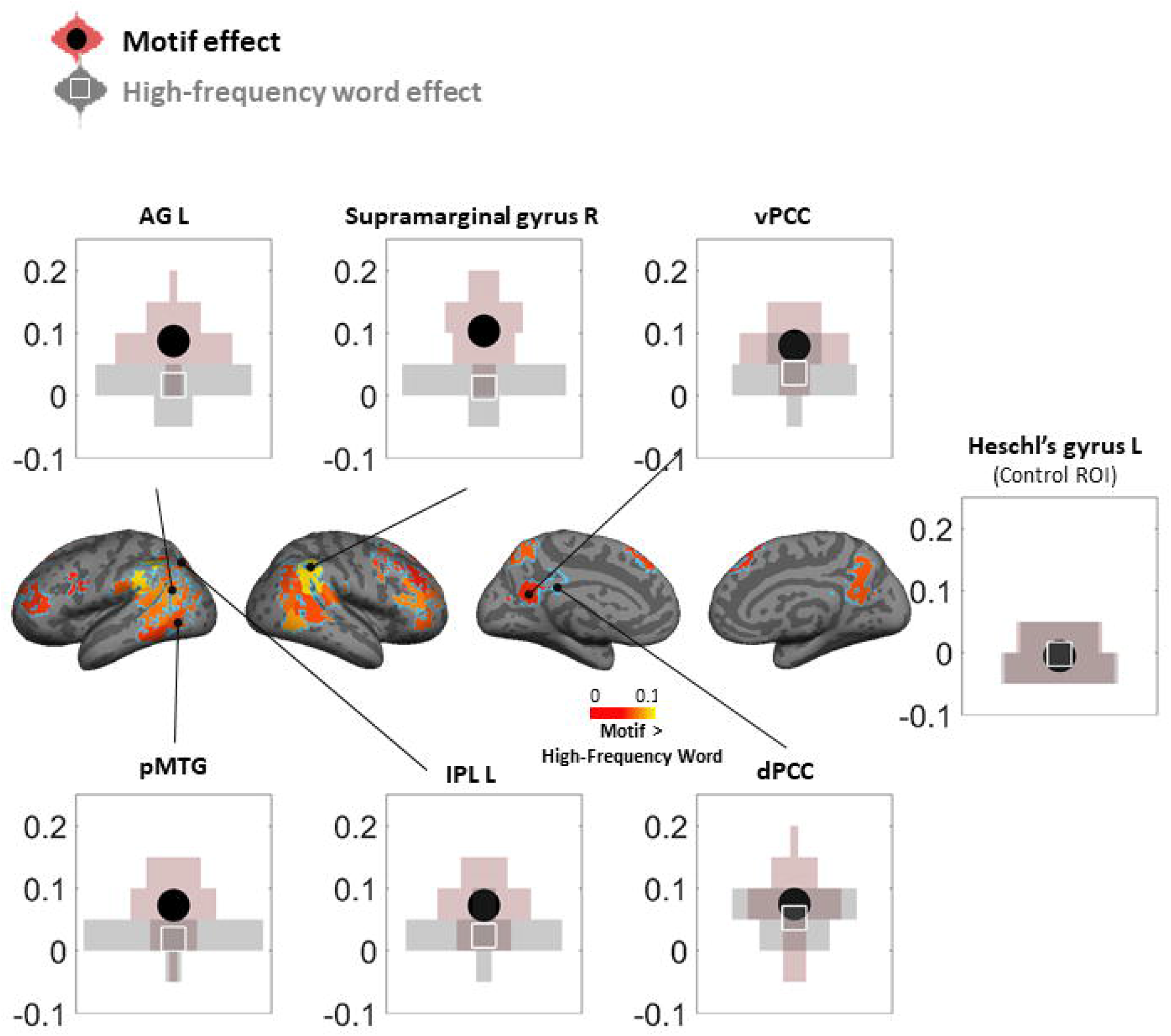
Motif vs. high-frequency word effect. Among regions showing a significant motif effect (marked by the blue outline), the effect of storyline-specific high-frequency words was computed using the same RSA method. The two effects were compared using one-tailed one-sample t-test (N=25, p < .05, FWE). Shaded areas show the distribution across subjects.

### Correlation between motif reinstatement and the behavioral relation score

To test whether participants showing stronger neural reinstatement also did a better job of integrating related events, we evaluated how well participants understood the relation between the AB and C events using 14 open questions in the post-scan questionnaire. The fMRI participants’ answers were evaluated by 5 raters who were blind to our hypothesis (3 females, age 26-31). The raters were asked to judge whether the fMRI participants responded by remembering only the AB part (score 0), only the C part (score 0), or both (score 1). Responses indicating no memory or false memory were given a score of 0.

Below are two sample questions and real responses from the fMRI participants (see Supplementary Table 4 for all the answers and scores):

*The following prompts are words, sentences, or phrases that recurred in the story. Please explain their significance to the story:*

*Q. Homemade chili=?*

*Ans. (average score 1): Clara makes homemade chili in the beginning and we find out that Clara learned the recipe from Steven, but Steven makes it better.*

*Ans. (average score 0): Steven makes it.*

*Ans. (average score 0): what Clara makes and Gary forgets to eat*

*Q. Mustard stained blouse =*

*Ans. (average score 1): Margaret stains her blouse with mustard when meeting with Alexander, also Steven can’t bear to wash it after Margaret dies.*

*Ans. (average score 0): Margaret’s blouse is stained when she meets with Alexander for the first time*

*Ans. (average score 0): Clara eating hotdogs in NYC*

We correlated (across participants) the size of the neural motif reinstatement effect for each participant with the participant’s ability to relate the AB part of the story to the C part, as revealed by their average relation score. This analysis was run separately for each ROI that showed a significant motif reinstatement effect. Within these ROIs, we found a significant correlation between the neural reinstatement of motifs and the individual relation scores in left inferior parietal lobule (IPL), left supramarginal gyrus (SMG), left angular gyrus (AG), and right middle frontal gyrus (MFG) (one-tailed, FDR corrected q < .05, Figure 8). In other words, participants who showed a stronger neural reinstatement of motifs were also better at reporting the narrative related connections among separate events sharing the same motifs.

**Figure 8.**
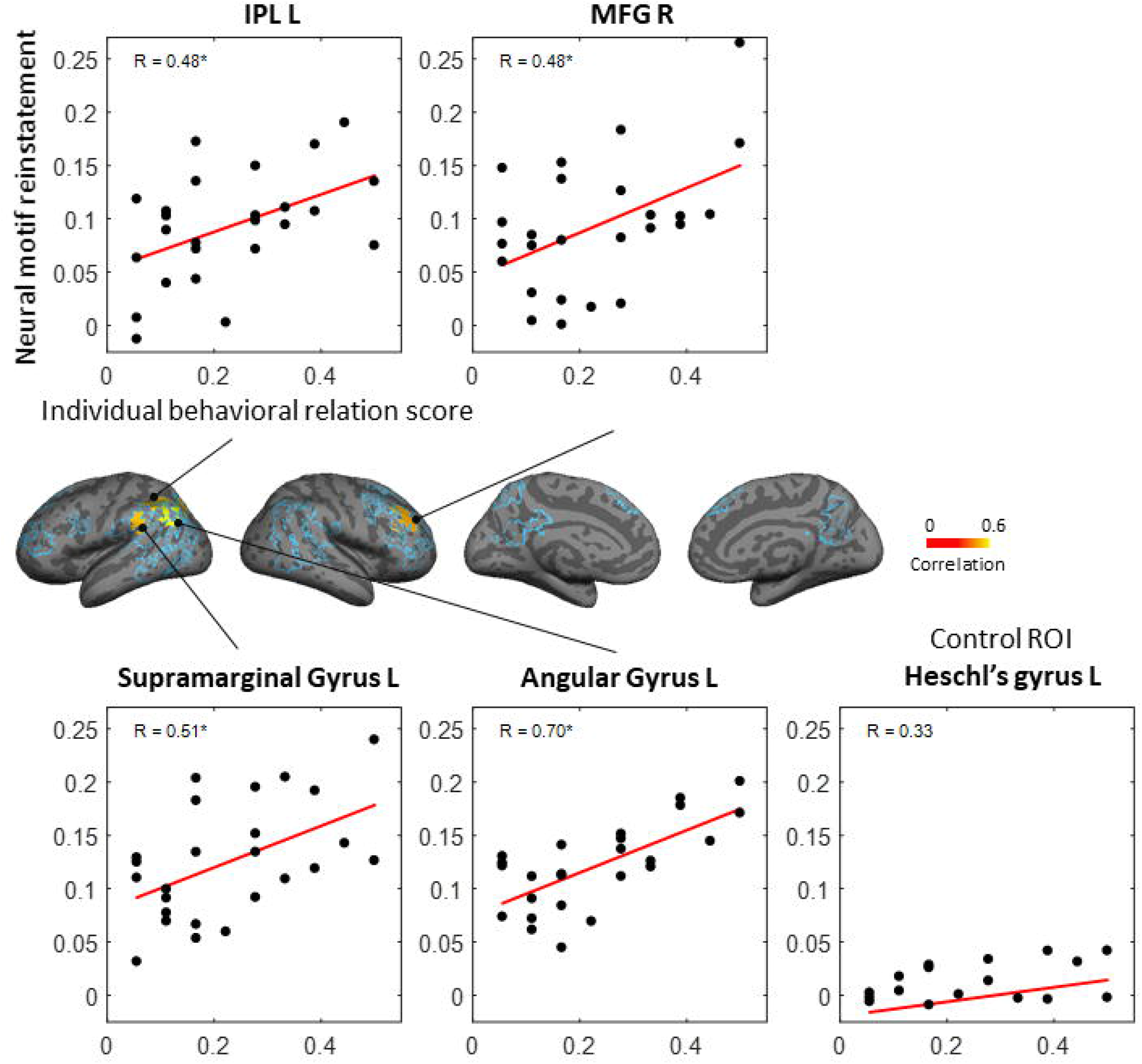
Correlation between the neural reinstatement of motifs and the individual relation scores. Each dot represents one subject. Only ROIs showing significant motif effect were tested (marked by the blue outline). Asterisks indicate significant correlation (N = 25, one-tailed, FDR corrected q < .05).

A possible alternative explanation of this result is that the relation score is tapping memory strength (not relations *per se*) and that the correlation between neural motif reinstatement and relation score simply reflects a shared influence of memory strength on these variables (i.e., high memory strength is associated with strong neural reinstatement and a high relation score, leading to a correlation between these two measures). To address this potential confound, we generated a separate “AB memory score” based on 14 questions in the post-scan questionnaire that only required a good memory of the AB part to answer (e.g. what does Clara find out during the party?); the mean level of accuracy for these AB memory score questions was 75% (SD: 17%). We then computed the partial correlation between the relation score and the neural motif effect, controlling for the AB memory score. The partial correlation effect was significant in left AG (R = 0.67, p < .001). Left SMG (R = 0.49, p = .008), left IPL (R = 0.42, p = .020), and right MFG (R = 0.42, p = .020) also showed partial correlations that were (individually) significant, although they were not significant after FDR correction for multiple ROIs. The fact that the relation score and neural motif effect were still correlated, even after controlling for the AB memory score, indicates that this relationship can not merely be explained in terms of memory strength.

### Between-subject vs. within-subject RSA

We used between-subject RSA for the analyses described above because our previous study (Chen et al., 2017) found that neural patterns associated with the perception and retrieval of specific events in a movie are shared across subjects (for similar findings, see Baldassano et al., 2017, 2018; Zadbood et al., 2017) – to the extent that these patterns are shared, this suggests that averaging the neural patterns across subjects will boost the signal-to-noise ratio (SNR). For a similar argument and detailed analysis of such effects, see Simony et al. (2016). For completeness, we have also included within-subject versions of our analyses. As we predicted (based on our prior work), the results of these within-subject analyses are qualitatively similar to the across-subject analyses but somewhat weaker, presumably due to lower SNR (Supplementary Figure 5-9).

### Hippocampal-cortical ISFC and cortical reinstatement of storyline and motif

To examine whether storyline reinstatement is dependent on connectivity with the hippocampus, we examined the correlation between storyline effect and hippocampal-cortical inter-subject functional correlation (ISFC) (Simony et al., 2016) in ROIs showing a significant storyline effect. For each A/B segment (except for the first segment of each storyline), we correlated the storyline effect with hippocampal-cortical ISFC during that segment and ran the correlation across segments (within participants) and participants (within segments). Because we did not have strong predictions about the relevant time windows for computing the storyline effect and ISFC, we ran an exploratory grid search across a range of analysis parameters. The ISFC between hippocampus and mPFC showed a strong correlation with the storyline effect (FDR corrected q < .05 across ROIs) for multiple settings of analysis parameters (Figure 9), although the result did not survive multiple-comparisons correction when factoring in the full set of analysis parameters, so it should be interpreted with caution. We also examined the correlation between hippocampal ISFC and motif reinstatement in ROIs showing a significant motif effect but did not find a significant correlation.

**Figure 9.**
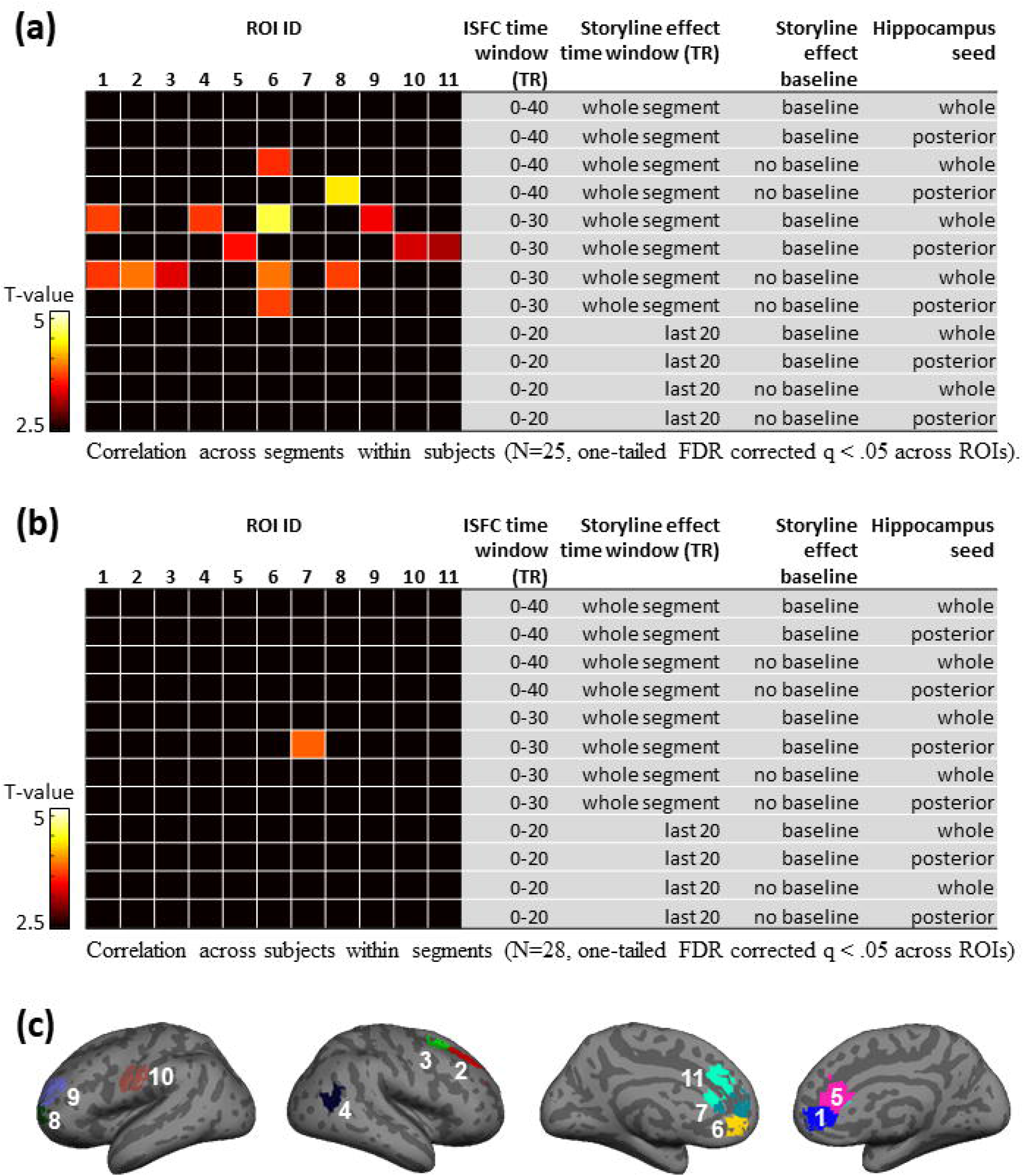
Correlation between hippocampal-cortical ISFC and storyline reinstatement within ROIs showing a significant storyline effect. (a) The correlation across segments was computed for each subject, excluding the first segment of each storyline. The resulting R-values were entered into a one-tailed t-test after Fisher’s z-transformation (FDR corrected q < .05 across ROIs). (b) Correlation across subjects was tested using the same methods. (c) Regions showing significant correlation in at least one of the analyses.

## Discussion

For this study, we actively designed a structured narrative in collaboration with a professional author to test how related events are dynamically and flexibly integrated by the brain while being protected and segregated from intervening irrelevant events. Our results indicate that the memory traces of recent events can be reactivated as a function of current input. This is seen in Figure 2b, where the neural patterns associated with the current storyline were reactivated at segment boundaries, while the activation patterns of the irrelevant storylines subsided. This effect was stronger in areas with longer processing timescales, peaking in the DMN (Figure 2a and 2c). The reinstatement of relevant past events was also tested in part C by using motifs to reactivate and update particular moments from both storylines (Figure 6). As predicted, the presentation of specific motifs in part C triggered the reinstatement of associated A/B events. Taken together, these results revealed a dynamic shift between currently active context and latent inactive contexts, which helps to integrate information over minute-long interruptions while protecting the accumulated information from irrelevant input.

Each storyline/scene is a unique combination of multiple narrative elements, such as characters, locations, goals, etc. In a prior paper (Yeshurun et al, PNAS, 2017), we showed that local differences in narrative elements (e.g. switching 2-3 words in a sentence with their antonyms) are amplified in the DMN, which showed response differences that are robust to spatial and temporal blurring. Furthermore, our prior work has shown that at least 15 dimensions of information are encoded in DMN activation patterns shared across subjects (Sherlock, Chen et al., 2017). Therefore, we speculate that the difference between the neural representations of the two storylines was driven by the unique combinations of narrative features, and should not be attributed to any single narrative dimension in isolation.

Using an audiovisual movie with interleaved storylines, a recent study (Milivojevic et al., 2016) reported the emergence of storyline-specific patterns in the hippocampus. However, as noted earlier, storyline information in that study was confounded with sensory features. Milivojevic et al. (2016) sought to control for these differences by regressing out sensory features, but this strategy is not ideal: When storyline and sensory information are strongly correlated, regressing out sensory information can attenuate legitimate storyline effects. Our study avoided this confound by presenting both storylines auditorily by the same speaker. We not only replicated the finding of neural differentiation in the hippocampus (Figure 4) but also revealed similar patterns of results in extended cortical areas, including language areas, the default mode network, the executive network, high order visual areas, and subcortical areas. In another related study, Lahnakoski et al. (2017) interleaved two independent movies and found that breaking the flow of a narrative (by interleaving) disrupts the accumulation of information relative to continuous viewing, similar to our previous studies (J. Chen et al., 2016; Honey, Thesen, et al., 2012; Lerner et al., 2011). However, unlike our study and Milivojevic et al. (2016), this study did not test for the coding of storyline-specific information and its reactivation across interruptions.

In line with the idea that the reactivated information is integrated with new input, we found that participants showing stronger neural reinstatement of motifs in left SMG, AG, IPL, and right MFG also showed higher relation scores (Figure 8). This finding supports the hypothesis that reinstatement of motifs led to better integration of AB and C parts. Also, we observed a storyline x time effect (Figure 3), which is consistent with the hypothesis that new segments updated the representations of the two storylines and pushed them further apart. It is worth noting that the storyline x time effect (computed using between-subjects RSA) does not on its own provide definitive evidence for the within-storyline integration. An alternative explanation for this result is that, over time, individual storylines become represented in a more stable and stereotyped fashion across listeners (thus making the two storylines more discriminable across listeners), as opposed to the storylines becoming more different *within* listeners. In principle, the within-subject RSA results (shown in Supplementary Figure 6) should be able to resolve this point, as any difference in the storyline effect over time in this analysis necessarily would reflect a within-subject effect. However, in practice, the results of this analysis are equivocal. The pattern of results across ROIs is qualitatively similar (suggesting that the storyline representations truly are moving apart) but the results no longer pass FDR correction; it is unclear whether this simply reflects the lower SNR of within-subject (vs. between-subject) analysis or the absence of a true “storyline differentiation” effect.

Lastly, having demonstrated the reinstatement of relevant past information, we next addressed the contribution of hippocampus-based episodic memory to this reinstatement. Previous work by Chen et al. (2016) showed that hippocampal-cortical interaction helped participants to integrate information across movie segments separated by a 1-day break. In our exploratory analyses, we found that storyline reinstatement increased with functional connectivity to hippocampus in mPFC both across subjects and across segments (Figure 9), for several (but not all) parameter settings for the analysis. This finding fits with the idea that the hippocampus, which is known to be involved with the reinstatement of episodic memories over days, may also be involved with the reinstatement of recently accumulated memories over shorter lags (for additional evidence in support of this view, see, e.g., Ezzyat & Olson, 2008; Goodrich, Baer, Quent, & Yonelinas, 2019; Hannula & Ranganath, 2009; Olson, Ingred, Sledge, Marianna, & Anjan, 2006). Notably, the degree of inter-subject functional connectivity between hippocampus and cortex did not reliably predict neural reinstatement triggered by motifs in part C. One possible explanation is that the inter-subject functional connectivity method involves averaging over multiple time points; this may make it less useful for detecting brief reinstatement events triggered by motifs.

In addition to hippocampus-mediated episodic memory, there are other mechanisms that might contribute to reinstating storyline representations. For example, recent studies of working memory have shown that past information could be held in the cortex during a delay period without persistent activity (Sprague, Ester, & Serences, 2016; Stokes, 2015; Wolff, Jochim, Akyürek, & Stokes, 2017) and such inactive neural patterns can be reactivated on task demand (Watanabe & Funahashi, 2014), by probe stimuli (Wolff et al., 2017), and by transcranial magnetic stimulation (Rose et al., 2016); several computational models have been built to account for the latent memory (Buonomano & Maass, 2009) and short-term synaptic plasticity in cortex has been proposed to be the underlying mechanism (Miller, Lundqvist, & Bastos, 2018; Mongillo, Barak, & Tsodyks, 2008; Zucker & Regehr, 2002). More research is needed to directly test whether short-term plasticity within cortex plays a role in supporting the memory of recent context.

Our design successfully induced reactivation of neural patterns associated with specific storylines and motifs, which, we believe, reflects the reinstatement of narrative information relating to past events rather than simple reactivation of word representations. In support of this view, we found a significant motif effect even when taking storyline-specific high-frequency words as the baseline (Figure 7), showing that word repetition is not sufficient to drive the effect. Also, the same motif was not always expressed in the same words (“throwing up” vs. “Clara feels sick, as the coffeecake rises to her throat”), showing that word repetition is not necessary to drive the effect. Furthermore, as noted above, we found a correlation between the behavioral relation scores and the neural motif effects (Figure 8). In other words, the same set of narrative motifs yielded greater neural reinstatement in subjects who demonstrated a better understanding of the narrative. As for the storyline effect, the strongest difference between storylines was found in regions with long TRW, i.e. regions where randomly ordered words do not elicit reliable responses (Lerner et al., 2011) (Figure 2c). Furthermore, 27% of the word tokens in the early AB part are storyline specific, while only 24% of the word tokens in the late AB part are storyline specific. Therefore, the difference in wording can not explain the finding of a stronger storyline effect in the later AB part (Figure 3).

In conclusion, real-life events require dynamic integration of past and present information. Our results suggest that process-memory may have two states, a state in which prior events are active and influence ongoing information processing, and an inactive state, in which the latent memory does not interfere with the ongoing neural dynamics (Hasson et al., 2015; Stokes, 2015). Through cross-disciplinary collaboration, this study demonstrated a way to achieve some experimental control over naturalistic stimuli and showed how skilled storytellers leverage these mechanisms of separation and integration to bring about the desired effects in the listener’s brain.

## Data availability

The data used in this study have been publicly released as part of the “Narratives” collection. Raw MRI data are formatted according to the Brain Imaging Data Structure (BIDS) with exhaustive metadata and are publicly available on OpenNeuro: https://openneuro.org/datasets/ds002245. The data corresponding to this study are indicated using the “21st year” task label. These data can be cited using the following reference: Nastase, S. A., Liu, Y.-F., Hillman, H., Zadbood, A., Hasenfratz, L., Keshavarzian, N., Chen, J., Honey, C. J., Yeshurun, Y., Regev, M., Nguyen, M., Chang, C. H. C., Baldassano, C. B., Lositsky, O., Simony, E., Chow, M. A., Leong, Y. C., Brooks, P. P., Micciche, E., Choe, G., Goldstein, A., Halchenko, Y. O., Norman, K. A., & Hasson, U. Narratives: fMRI data for evaluating models of naturalistic language comprehension. https://doi.org/10.18112/openneuro.ds002245.v1.0.3

## Acknowledgment

This study is supported by a Magic Grant from Princeton’s Humanities Council, Taiwan Ministry of Science and Technology (105-2917-I-564-007), and by the National Institute of Mental Health (R01-MH112357).

## Author contributions

C.L., Y.Y., and U.H. designed the experiment; C.L authored the stimulus; C.L and Y.Y created the audio rendition of the narrative; C.H.C. and Y.Y. acquired the data; C.H.C. analyzed and interpreted the data with input from U.H., C.L., and K.A.N.; C.H.C. drafted the manuscript; U.H., K.A.N., Y.Y., and C.L. substantively revised the manuscript.

## Additional information

### Competing financial interests

The authors declare no competing financial interests.

## Notes

### Competing Interest Statement

The authors have declared no competing interest.

