## Supplementary Figure 1-9 for "Relating the past with the present: Information integration and segregation during ongoing narrative processing"

#### (a) Audio Amplitude

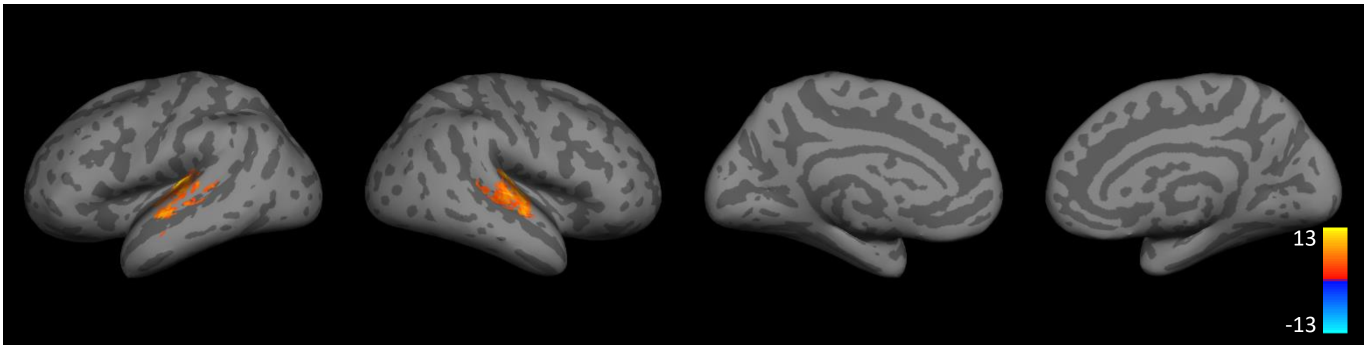


#### (b) Between-Segment Pause

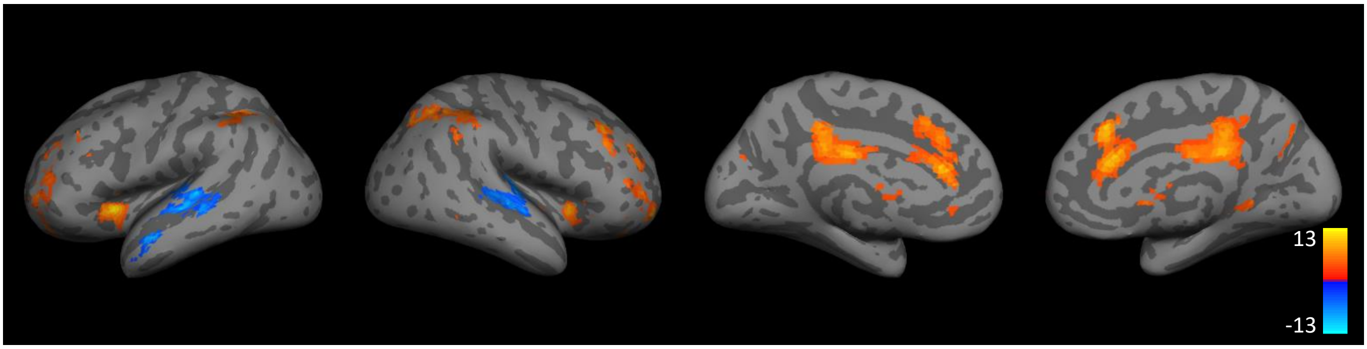


Supplementary Figure 1. T-value maps showing the effects of audio amplitude and between-segment pause (N=25, p < .05, FWE corrected).

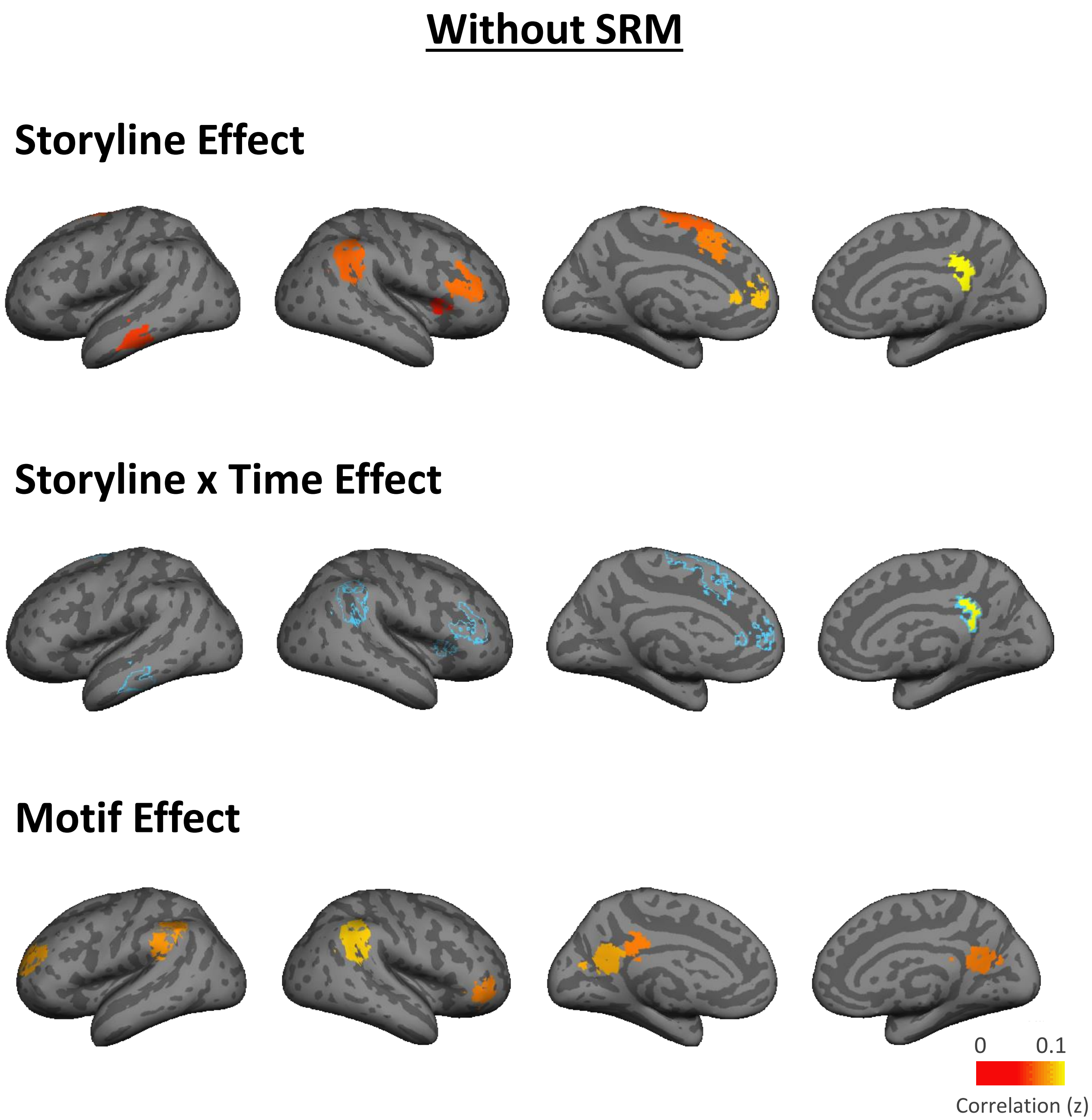


Supplementary Figure 2. Storyline and motif effects without applying SRM (p < .05. FWE)

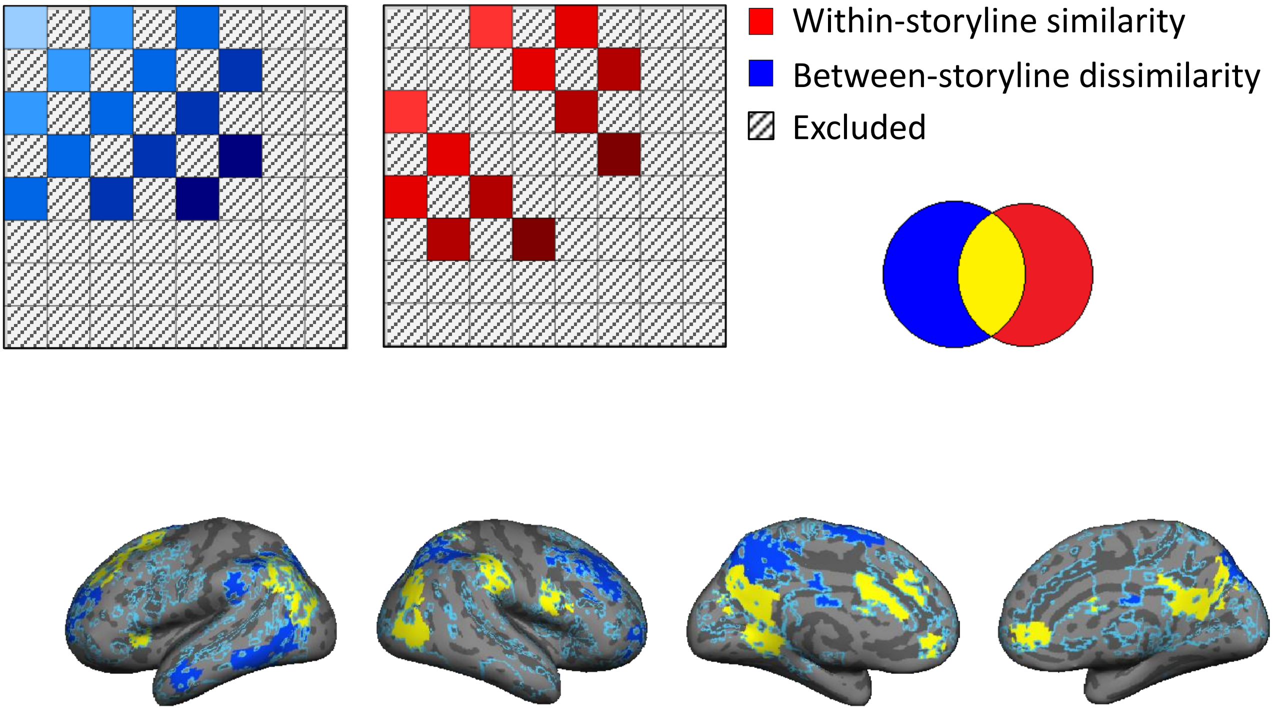


Supplementary Figure 3. Graded time x storyline effect. The time x between-storyline dissimilarity and time x within-storyline similarity effects were tested separately, within regions showing significant storyline effect (marked by blue outline) (N=25, $p < .05$, FWE) and their overlap is shown in the lower panel (yellow regions).

#### (a) Predicted Character Effect

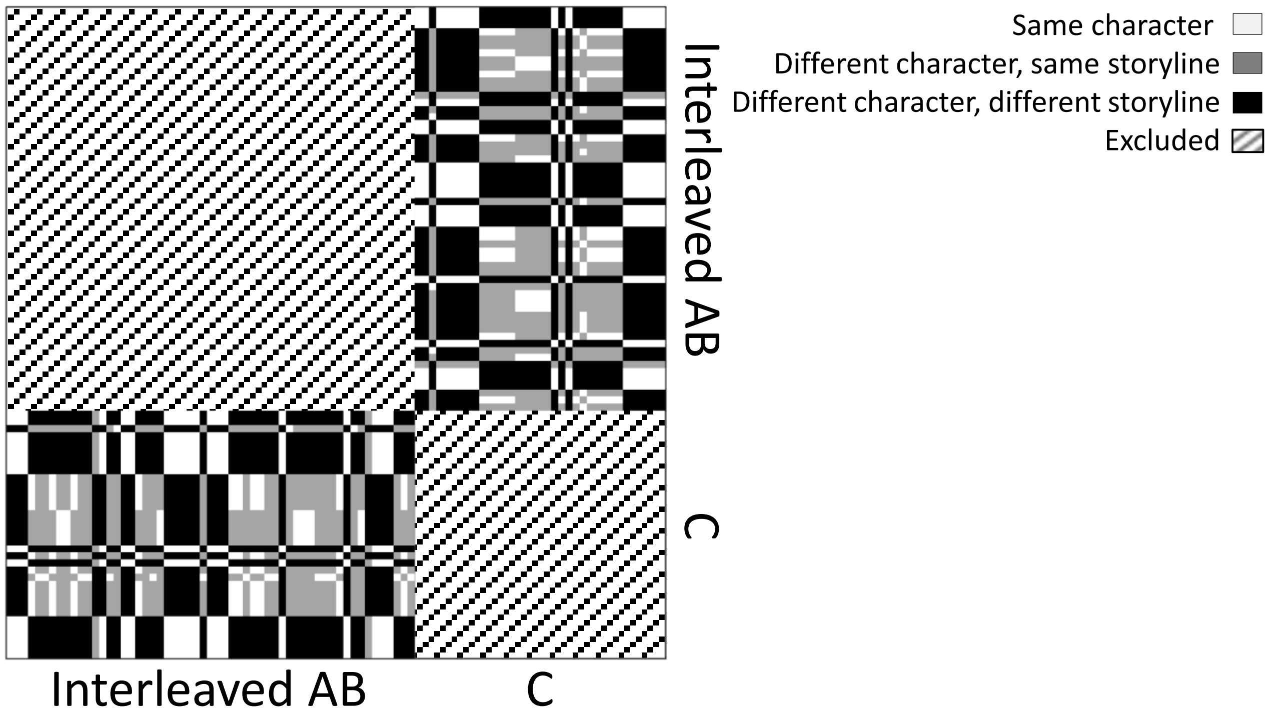


#### (b) Character Effect

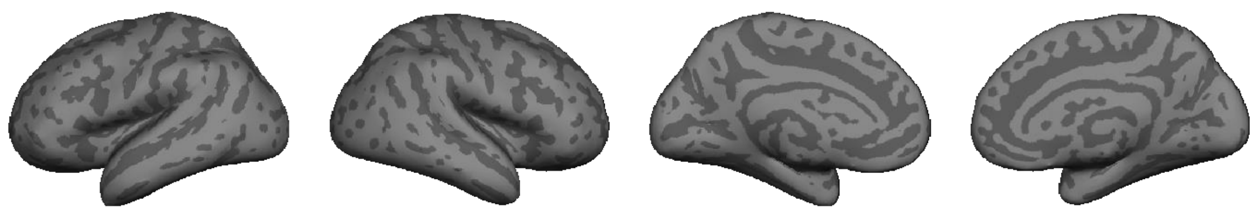

Supplementary Figure 4. (a) Predicted pattern similarity between motif-related events based on whether they share the same character. (b) character effect was not significant ($p < .05$, FWE).

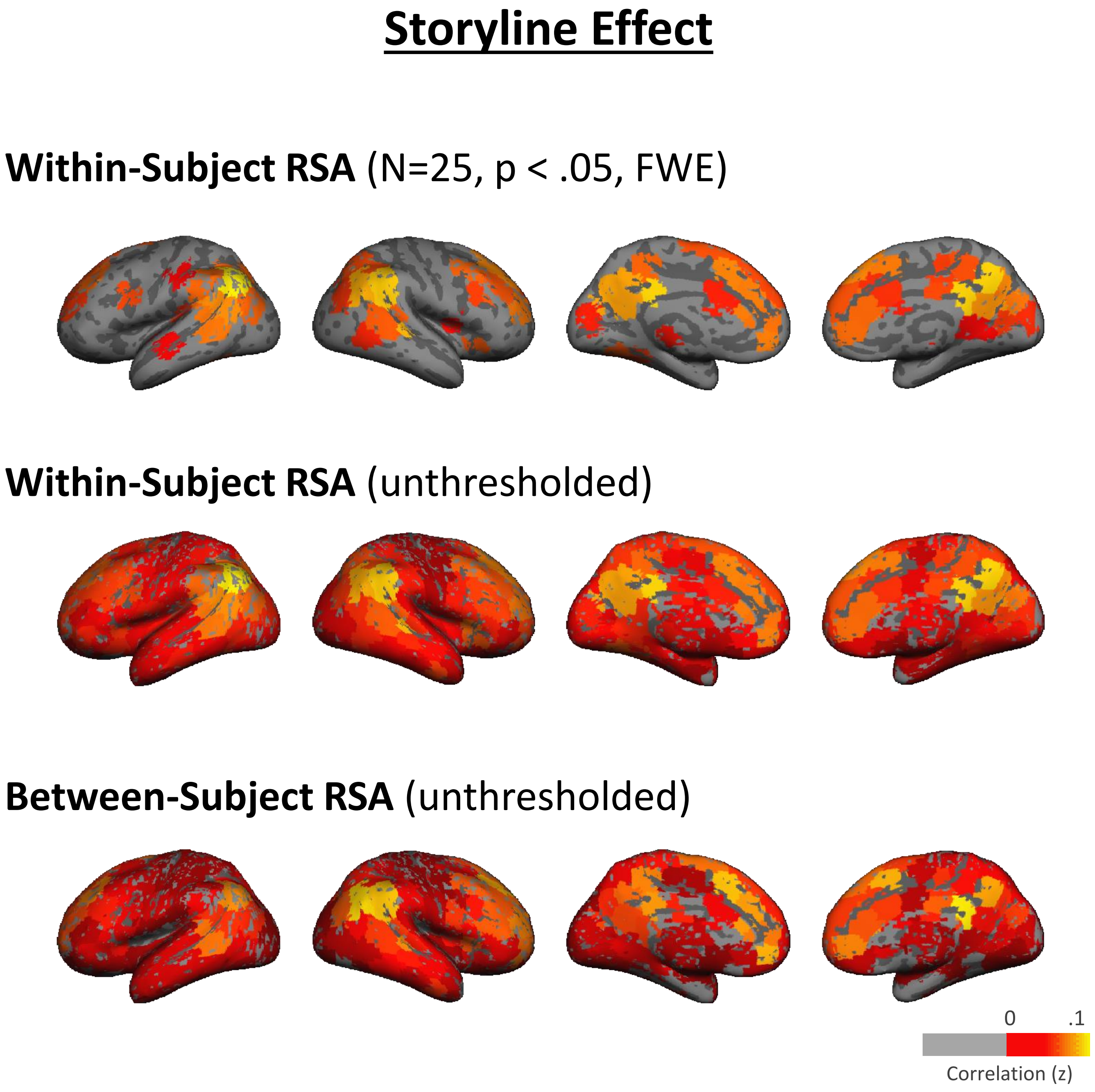


Supplementary Figure 5. Comparison between the within-subject RSA and between-subject RSA of the storyline effect. For the thresholded result of between-subject RSA, see Figure 2.

### Storyline x Time Effect

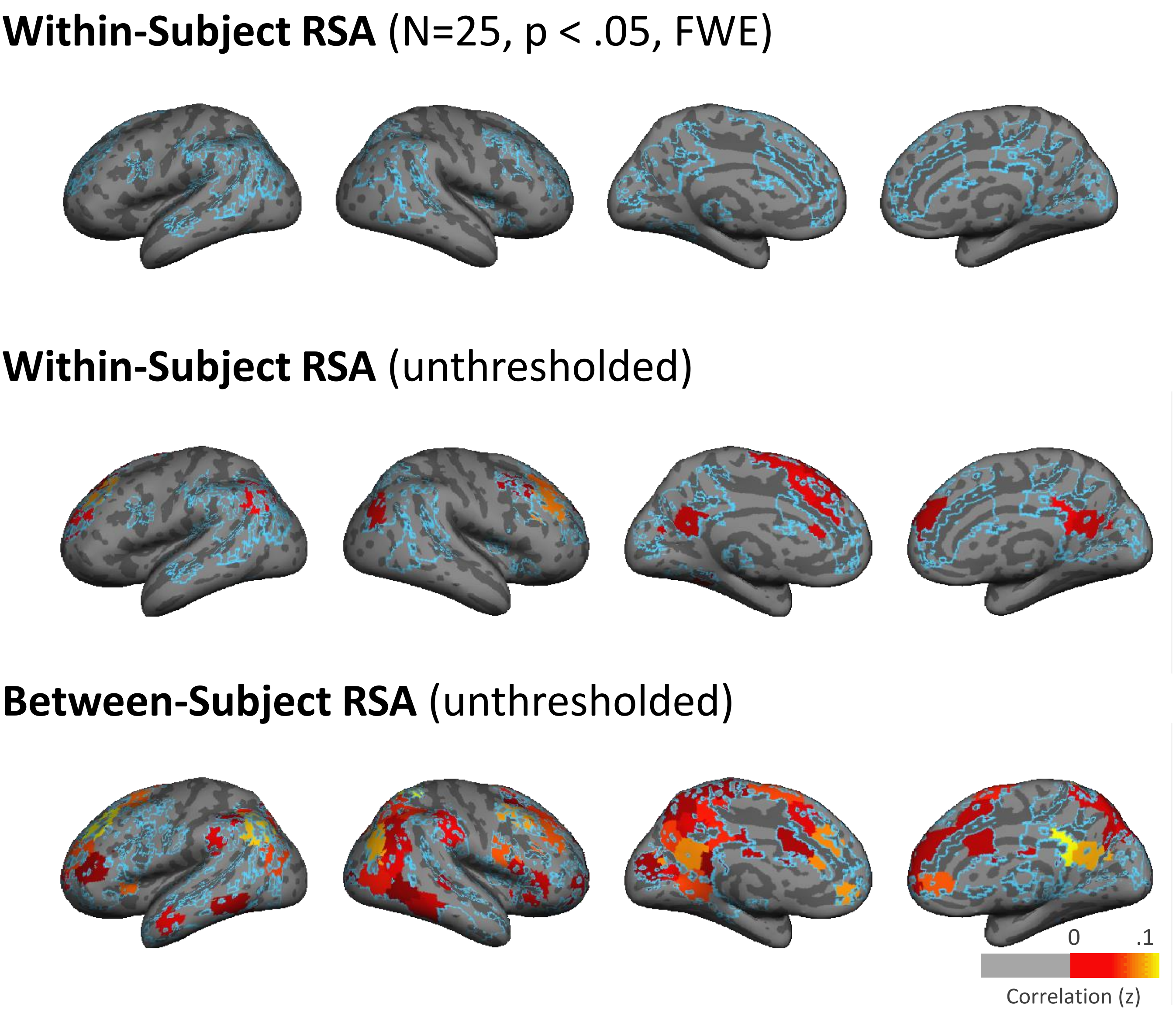


Supplementary Figure 6. Comparison between the within-subject RSA and between-subject RSA of the storyline x time effect. Only ROIs showing significant storyline effect were tested (marked by the blue outline). For the thresholded result of between-subject RSA, please see Figure 3.

### Motif Effect

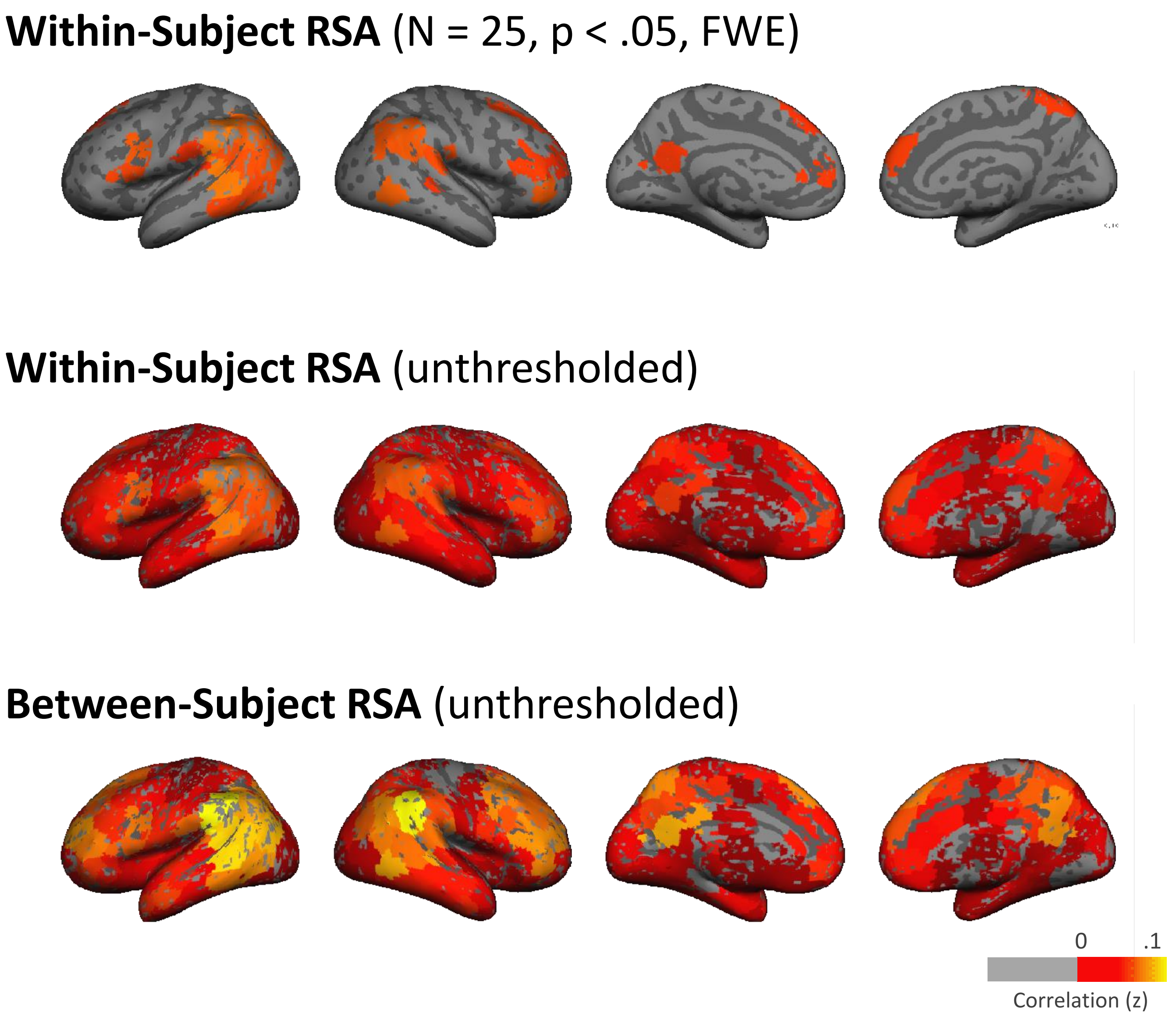


Supplementary Figure 7. Comparison between the within-subject RSA and between-subject RSA of the motif effect. For the thresholded result of between-subject RSA, please see Figure 6.

### Correlation Between Motif Effect and Relation Score

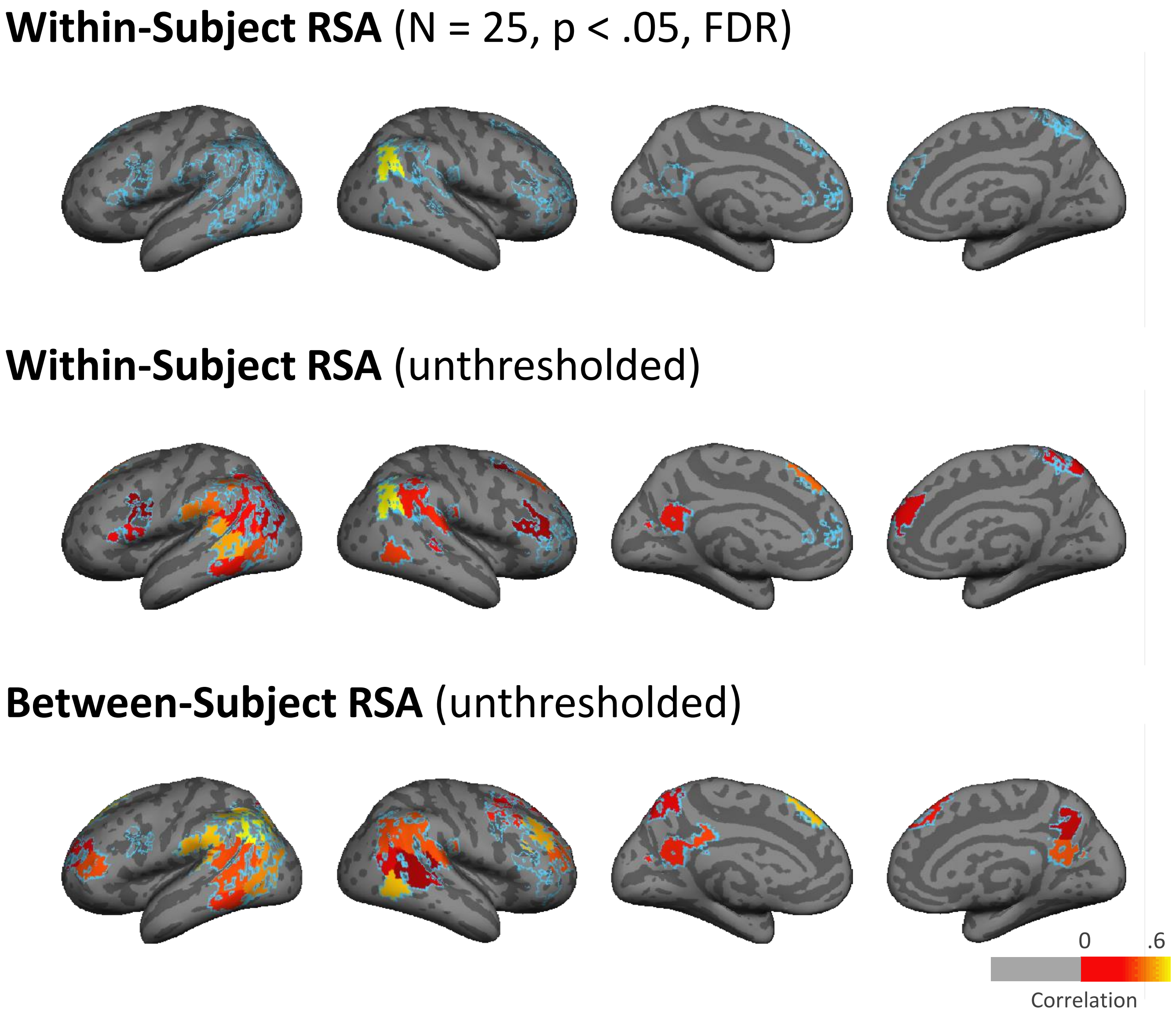


Supplementary Figure 8. Correlation between the relation score and motif effect computed using within-subject vs. between-subject RSA. Only ROIs showing significant motif effect were tested (marked by the blue outline). For the thresholded result of between-subject RSA, please see Figure 8.

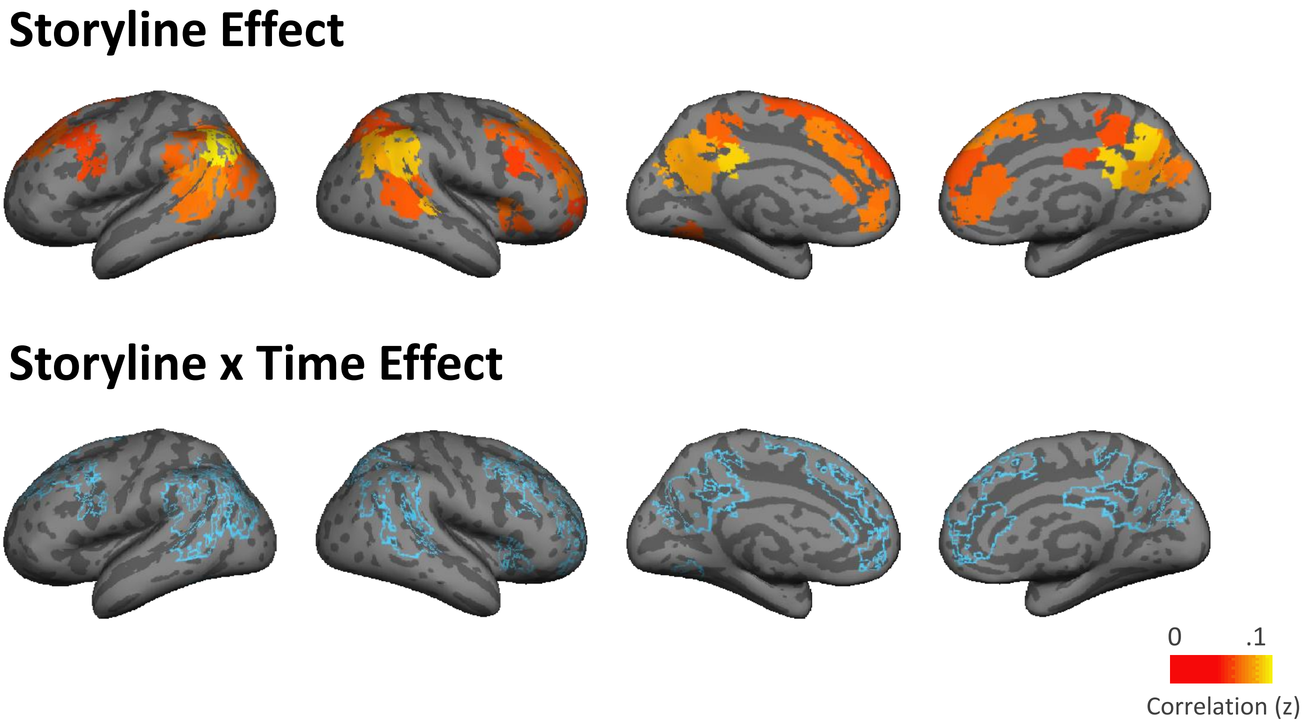


Supplementary Figure 9. Within-subject RSA of the storyline effect and storyline x time effects, statistical threshold obtained by shuffling the labels of A/B segments ($p < .05$. FWE) instead of group t-test.
