## Supplementary Table 3 for "Relating the past with the present: Information integration and segregation during ongoing narrative processing"

Supplementary Table 3. Storyline specific high-frequency words used for the control analysis (Figure 7).

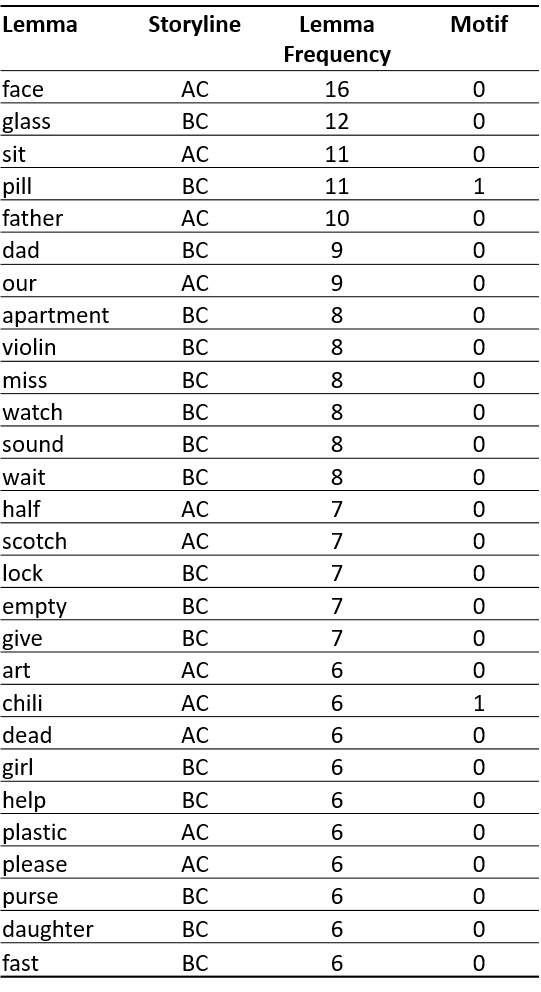
